# Serine-ubiquitination regulates Golgi morphology and the secretory pathway upon Legionella infection

**DOI:** 10.1101/2020.07.27.223842

**Authors:** Yaobin Liu, Rukmini Mukherjee, Florian Bonn, Thomas Colby, Ivan Matic, Marius Glogger, Mike Heilemann, Ivan Dikic

## Abstract

SidE family of *Legionella* effectors catalyze non-canonical phosphoribosyl-linked ubiquitination (PR-ubiquitination) of host proteins during bacterial infection. SdeA localizes predominantly to ER and partially to the Golgi apparatus, and mediates serine ubiquitination of multiple ER and Golgi proteins. Here we show that SdeA causes disruption of Golgi integrity due to its ubiquitin ligase activity. The Golgi linking proteins GRASP55 and GRASP65 are PR-ubiquitinated on multiple serine residues, thus preventing their ability to cluster and form oligomeric structures. In addition, we found that the functional consequence of Golgi disruption is not linked to the recruitment of Golgi membranes to the growing *Legionella*-containing vacuoles. Instead, it affects the secretory pathway, including cytokine release in cells. Taken together, our study sheds light on the Golgi manipulation strategy by which *Legionella* hijacks the secretory pathway and promotes bacterial infection.

## Introduction

Ubiquitination is a post-translational modification that is conserved from yeast to mammals. The catalysis of canonical ubiquitination is regulated via a three-enzyme cascade: firstly an ubiquitin (Ub) molecule is activated by an Ub-activating enzyme (E1) utilizing ATP; the activated Ub is linked to the catalytic cysteine of the E1 via its C-terminal Glycine and subsequently transferred to an Ub-conjugating enzyme (E2); finally, a Ub ligase (E3) links the carboxyl group of the ubiquitin’s C-terminal glycine to the ε-amino group of the target lysine in the substrate protein by an isopeptide bond (Hershko et al., 2000). Protein ubiquitination virtually regulates every cellular processes, including protein quality control, protein trafficking, immunity, and DNA repair by targeting substrates to the proteasome or altering their functions (Ben-Neriah, 2002; Dikic, 2017; Donaldson et al., 2003; Rape, 2018).

Consistent with the critical roles of ubiquitination in cellular processes, emerging evidence indicates that pathogens hijack the ubiquitination machinery for efficient invasion (Bomberger et al., 2011; Hicks and Galán, 2013; Maculins et al., 2016). For example, the intracellular Gram-negative pathogen *Legionella pneumophila* secretes more than 300 effectors into host cells via its type IV secretion system (T4SS) (Hubber and Roy, 2010). Many of these toxins function as E3 ligases and are reported to manipulate host ubiquitination (Qiu and Luo, 2017). Various studies have revealed that effectors of the SidE family (SdeA, SdeB, SdeC and SidE) catalyze an NAD^+^-dependent, ATP-independent type of ubiquitination without the need of E2 and E3 enzymes (Bhogaraju et al., 2016; Qiu et al., 2016). Moreover, unlike the conventional ubiquitination that occurs on lysine residues of substrate proteins, SidE family effectors catalyze the conjugation of Ub via a phosphoribosyl moiety to serine residues of host substrate proteins by a two-domain catalytic relay: a mono ADP-ribosyl transferase (mART) domain that ADP-ribosylates Arg42 of Ub and a phosphodiesterase (PDE) domain that cleaves the phosphodiester bond of the ADP-ribosylated Ub (ADPR-Ub) and conjugates the resulting phosphoribosyl ubiquitin (PR-Ub) to the serine residue of a substrate (Akturk et al., 2018; Dong et al., 2018; Kalayil et al., 2018; Wang et al., 2018). PR-ubiquitination is reversible, and DupA is a deubiquitinase with specific affinity for PR-ubiquitinated substrates (Shin et al., 2020). SidE family effectors are crucial for bacterial virulence, as a *Legionella* strain lacking SidE family members shows defective growth in host cells, a phenotype that can be rescued by replenishment of SdeA (Bardill et al., 2005; Qiu et al., 2016). To date, numerous ER-associated proteins have been identified as PR-ubiquitination substrates of SdeA, such as tubular ER protein RTN4, FAM134B, and LNP1. PR-ubiquitination of these proteins is involved in regulating ER remodeling and recruiting ER membranes to *Legionella* containing vacuoles (LCV) where the bacteria resides and replicates (Kotewicz et al., 2017; Shin et al., 2020).

In our previous study, we used the catalytically dead mutant of the deubiquitinase DupA as a bait to identify targets of SdeA. Besides ER-related substrates, we also identified proteins related to other cellular pathways, including Golgi proteins, mitochondrial proteins and components of the autophagy machinery (Shin et al., 2020). However, the biological functions of PR-ubiquitination of these proteins remained unclear. In the present study, we made use of biochemical and microbiological approaches to characterize the PR-ubiquitination of Golgi tethering proteins GRASP55 and GRASP65 by SdeA. We also provide explanations for the Golgi morphological regulation by the PR-ubiquitination of these proteins. Moreover, we demonstrate that PR-ubiquitination regulates the host cellular secretory pathway during bacterial infection.

## Results

### SdeA is targeted to the ER and Golgi via its carboxyl terminus

Previous structural and biochemical studies have revealed the structure of SdeA catalytic core, and the mechanism by which SdeA ubiquitinates substrates is well established (Akturk et al., 2018; Dong et al., 2018; Kalayil et al., 2018; Wang et al., 2018). However, the function of the carboxyl terminal (CT) region, predicted to be coiled-coil, remained unknown (Fig. 1A). Previous reports suggested that coiled-coil domains are required for membrane localization of many *Salmonella* type III effectors (Knodler et al., 2011). In view of that SdeA co-localizes with ER protein calnexin and ubiquitinates many ER proteins, such as RTN4 and FAM134B (Kotewicz et al., 2017; Qiu et al., 2017; Shin et al., 2020), we hypothesized that the CT domain of SdeA is responsible for its membrane association. To analyze if the CT of SdeA contributes to its membrane localization and is therefore needed for the PR-ubiquitination of membrane-located substrates, we first investigated the ER localization of wild-type SdeA and truncated SdeA^1-972^ mutant lacking the last part of the C-terminal region (Fig. 1A). COS7 cells were transfected with plasmids encoding EGFP-tagged SdeA, a truncated mutant, or EGFP alone and subsequently stained for the ER resident protein Calnexin. We observed that ectopically expressed SdeA co-localized with ER protein Calnexin in COS7 cells, consistently with a previous study (Qiu et al., 2017). In contrast and along our hypothesis that the C-terminal region of SdeA is essential for its membrane localization, truncated SdeA did not co-localize with Calnexin but showed a rather cytosolic distribution similar to the EGFP control (Fig. 1B). In addition, we observed that part of SdeA was densely localized close to the nucleus in cells (Fig. 1B). Staining with the Golgi marker GM130 revealed that this part of SdeA co-localized with the Golgi apparatus, while the truncated mutant SdeA^1-972^ did not (Fig. 1C). We confirmed that the C-terminus region of SdeA is necessary to its Golgi localization by expressing the N-terminal-truncated SdeA^909-C^ in cells stained with GM130. SdeA^909-C^ expressed in cells highly overlapped with GM130, but did not disturb Golgi structure as wild-type SdeA did (Fig. 1C). This data suggests that the C-terminal part of SdeA is critical for its ER as well as its Golgi membrane localization.

**Figure 1.**
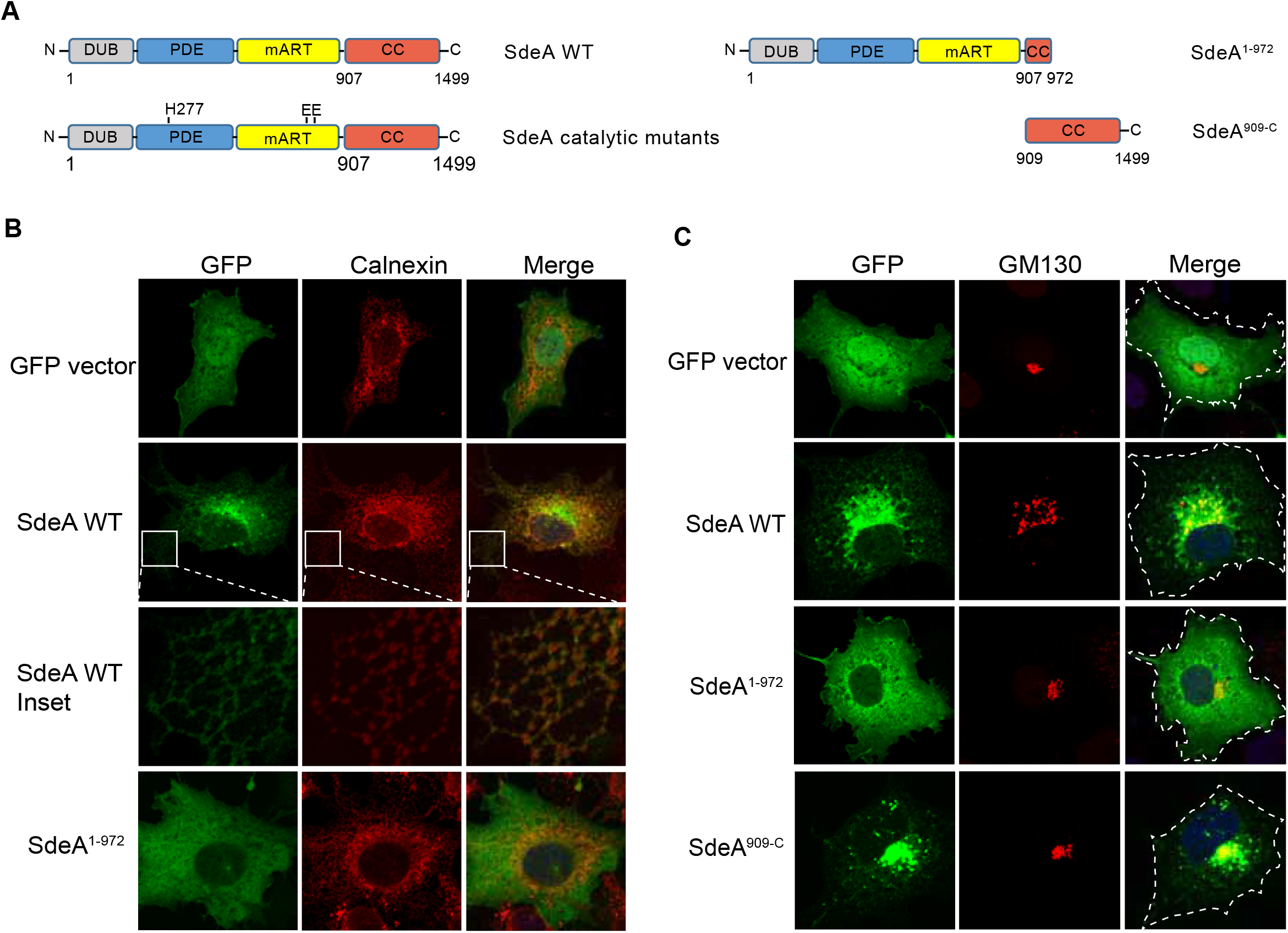
SdeA partially localizes to the Golgi. (**A**) Schematic diagrams of full-length wild-type SdeA, SdeA catalytic mutant SdeA H277A or SdeA EE/AA, truncated SdeA^1-972^ and SdeA^909-C^. (**B**) Confocal images showing the co-localization of SdeA (green) with ER protein Calnexin (red). COS7 cells were transfected with plasmids encoding GFP-tagged wild type SdeA or truncated mutant. Cells were cultured for 24 hours after transfection, then fixed, permeabilized, and stained with Calnexin antibody and visualized using confocal microscope. (**C**) Confocal images showing the co-localization of SdeA (green) with Golgi protein GM130 (red). Cells were cultured for 24 hours after transfection, then fixed, permeabilized, and ultimately stained with GM130 antibody and visualized using confocal microscope.

### SdeA induces disruption of Golgi structure

To investigate whether Golgi localization of SdeA is critical for its ligase function, we co-expressed wild-type SdeA or the truncated mutant SdeA^1-972^ with its known Golgi associated substrate Rab33b (Qiu et al., 2016). Western blot analysis showed that the truncated form of SdeA could not ubiquitinate Rab33b even though it was able to ADP-ribosylate ubiquitin (Figure 2-figure supplement 1A). This data suggests that the C-terminus region of SdeA is critical not only for its localization but also for its ability to ubiquitinate Golgi proteins. During our localization studies, we observed that expression of wild-type SdeA, but not the CT-truncated mutants, results in dispersed GM130 staining. This implicates an effect of SdeA activity on the structural organization of the Golgi apparatus. We then sought to investigate the possibility of PR-ubiquitination activity of SdeA regulates Golgi assembly by comparing the effects between wild-type SdeA with SdeA catalytic mutants (Fig. 1A). Expression of PDE defective mutant (SdeA H277A) or mART defective mutant (SdeA EE/AA) did not exhibit significant impact on the structure of the Golgi (Fig. 2A, B). In addition, the effect of wild-type SdeA on the Golgi structure could be counteracted by co-expression of DupA, the specific deubiquitinase for PR-ubiquitination, but not its catalytically dead mutant DupA H67A (Fig. 2A, B). These findings suggest that the Golgi disruption observed in cells expressing SdeA is likely to be caused by the accumulation of its ubiquitinated substrates. These observations are in apparent agreement with previous study (Jeong et al., 2015). Similar observations were also made in HeLa cells stained with both cis (GM130) and trans (TGN46) Golgi marker antibodies (Figure 2-figure supplement 1B). In order to evaluate the physiological relevance of SdeA in triggering Golgi disruption, we infected human lung carcinoma A549 cells with either a wild-type *Legionella* strain, a mutant strain missing genes encoding SidE family proteins (*ΔsidEs*) or a mutant that does not express DupA and DupB (*ΔdupA/B*). As expected, we observed a scattering of the Golgi apparatus in cells infected with wild-type but not *ΔsidEs Legionella* or control cells. Infection by *Legionella* without DupA/B caused more dramatic dispersal of the Golgi, compared to the wild-type *Legionella* (Fig. 2C, D). To further dissect the modulation of Golgi by SdeA, we analyzed the Golgi morphology of the cells expressing SdeA with super-resolution microscopy and electron microscopy. Data from DNA-PAINT super-resolution microscopy shows that cis Golgi protein GM130 and trans Golgi protein Golgin97 were still colocalized in cells expressing SdeA (Figure 2-figure supplement 1C), which is the same with the confocal image showing the colocalization of cis and trans Golgi proteins in HeLa cells expressing SdeA (Figure 2-figure supplement 1B). These results indicate that Golgi stacking may not be affected by SdeA. Moreover, SdeA expression did not change the level of the proteins required for Golgi structure maintenance (Figure 2-figure supplement 1D). Taken all together, these data suggest that SdeA mediated PR-ubiquitination of host substrates induces disruption but not complete fragmentation of Golgi ribbon.

**Figure 2.**
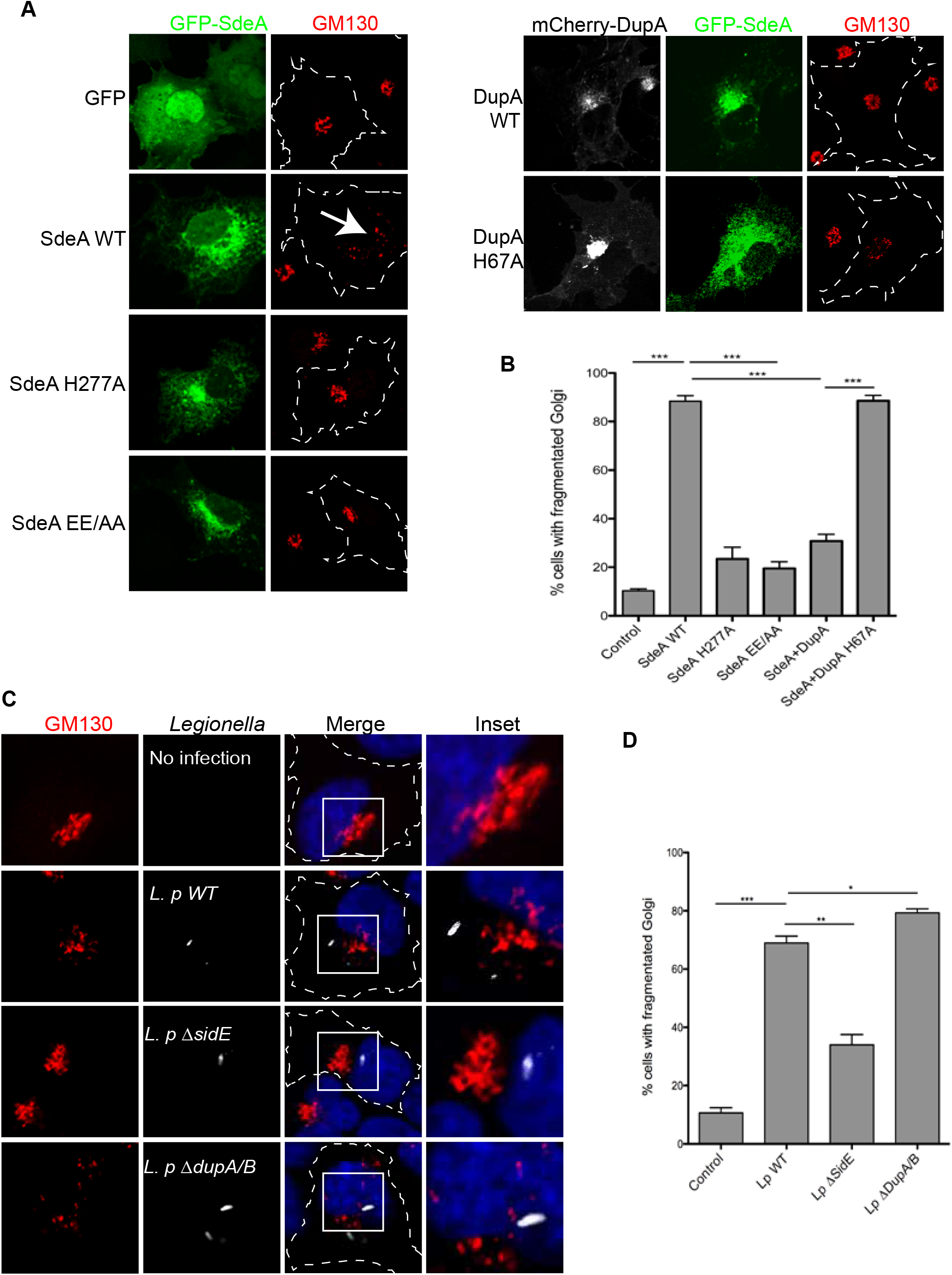
*Legionella* effector SdeA mediates Golgi fragmentation in cells. (**A**) Confocal images showing Golgi (red) fragmentation caused by exogenously expressed SdeA (green). GFP-tagged SdeA wild-type or catalytically defective mutants were expressed or co-expressed with DupA in COS7 cells. Cells were cultured for 24 hours after transfection then fixed with 4% PFA. (**B**) Quantification of the percentage of cells with dispersed Golgi in (**A**). Data are shown as means ± SEM of more than 60 cells taken from three independent experiments. ***P<0.001. (**C**) Confocal images showing Golgi fragmentation caused by *Legionella*. A549 cells were infected with wild-type or mutant *Legionella* as indicated. Cells were washed 3 times with PBS after 2 hours infection to remove non-phagocytosed bacteria, then fixed with 4% PFA and stained with indicated antibodies. (**D**) Quantification of the percentage of cells with dispersed Golgi in (**C**) Data are shown as means ± SEM of more than 70 cells taken from three independent experiments. Data were analyzed with unpaired t test, ***P<0.001, *P<0.01, *P<0.05.

### *In vitro* and *in vivo* validation of PR-ubiquitination of Golgi substrates by SdeA

Using the PR-deubiquitinase DupA as a bait, we pulled-down over 180 potential host substrate proteins of SdeA upon *Legionella* infection (Shin et al., 2020). Among these identified proteins, a number of ER resident proteins, and proteins related to Golgi components were highly enriched. Notably, Golgi proteins GRASP55 and GCP60 had the highest ratios among the putative Golgi protein substrates (Fig. 3A). Since GRASP55 play roles in the maintenance of the Golgi structure (Grond et al., Sohda et al., 2001), we hypothesized that SdeA modifies and inactivates Golgi proteins related to structure maintenance, thereby inducing Golgi disruption. GRASP65, which shares high sequence similarity with GRASP55, is localized to the *cis* Golgi and is also found in dispersed Golgi apparatus in cells expressing wild-type SdeA (Figure 3-figure supplement 1A). *In vitro* ubiquitination assays were performed, incubating purified GRASP55 or GRASP65 with SdeA and ubiquitin for 30 min, to monitor potential PR-ubiquitination of these two Golgi proteins. We observed that SdeA is able to modify both GRASP55 and GRASP65 *in vitro* (Fig. 3B). Furthermore, cellular expression of wild-type SdeA, but not inactive PDE or mART mutants, resulted in the appearance of ubiquitinated GRASP55 and GRASP65. This PR-ubiquitination was lost when PR-ubiquitination specific deubiquitinase, DupA, was co-expressed with wild-type SdeA (Fig. 3C, D). Similar observations were made for GCP60, where purified GCP60 from cells incubated with wild-type SdeA exhibited PR-ubiquitination (Figure 3-figure supplement 1B). Such modification also appeared in cells when GCP60 was co-expressed with wild-type SdeA but not upon co-expression of SdeA EE/AA mutant (Figure 3-figure supplement 1C). Along our hypothesis that SdeA is actively targeted to the Golgi, exogenous expression of CT-truncated SdeA mutants showed markedly reduced activity in modifying substrate GRASP55 (Figure 3-figure supplement 1D), similar to the effect observed on PR-ubiquitination of Rab33b, indicating that Golgi localization of SdeA is important for substrate modification. Our previous work indicated that M408 and L411 are two essential amino acids in the substrate binding region of SdeA (Kalayil et al., 2018). To distinguish whether SdeA targets GRASP55 protein specifically via its substrate recognition region or if modification is an overexpression artifact and due to high amounts of SdeA located at the Golgi, we performed an *in vitro* ubiquitination assay by incubating purified GRASP55 with wild-type SdeA and SdeA ML/AA mutant, respectively. The Coomassie staining showed that the SdeA ML/AA mutant did not ubiquitinate GRASP55 (Figure 3-figure supplement 1E). Similarly, GRASP55 ubiquitination is reduced in cells expressing SdeA ML/AA mutant compared to cells expressing wild type SdeA (Figure 3-figure supplement 1F) and the interaction with GRASP55 is much reduced for SdeA mutant compared to wild-type SdeA in Co-IP experiments (Figure 3-figure supplement 1G). Together, these results suggest that the PR-ubiquitination of Golgi tethering proteins GRASP55, GRASP65 and GCP60 by SdeA is a selective and functional part of the hijacking strategy of *Legionella*.

**Figure 3.**
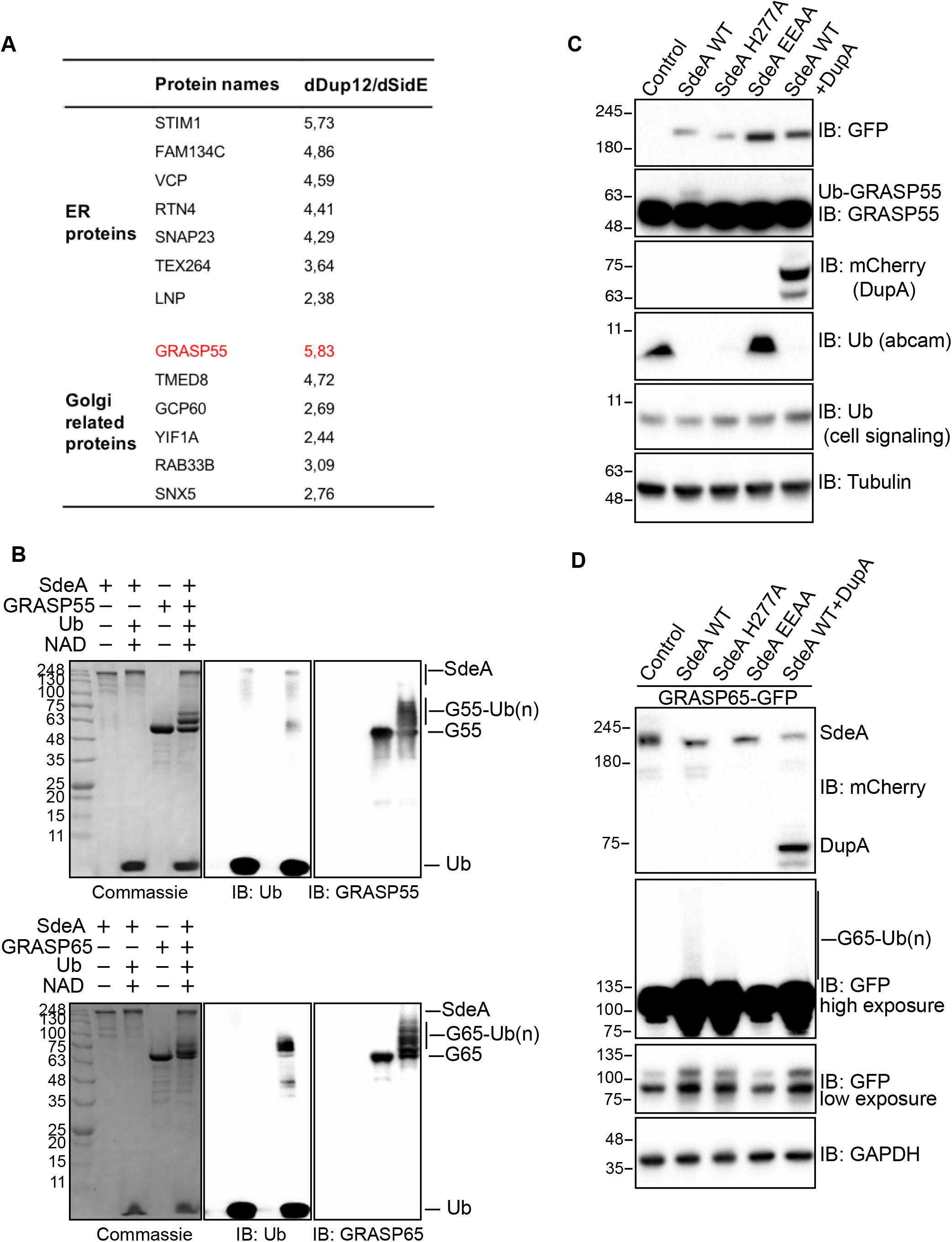
SdeA ubiquitinates Golgi tethering factor GRASP proteins. (**A**) Potential ER and Golgi protein substrates of SdeA identified by mass spectrometry. Values indicate intensity ratios between proteins enriched from samples infected with different *Legionella* strains (*ΔdupA/B* over *ΔsidE*). Among the substrate candidates, Golgi tethering factor GRASP55 (red) is one of the highly ubiquitinated proteins. (**B**) GRASP55 and GRASP65 ubiquitination by SdeA *in vitro*. Purified GRASP55 or GRASP65 were incubated with SdeA in the presence of NAD^+^ and ubiquitin. Reaction products were separated with SDS-PAGE and then stained with Coomassie blue or blotted with antibodies against ubiquitin, GRASP55 or GRASP65. (**C**) Modification of GRASP55 by exogenous SdeA in cells. HEK293T cells were transfected with wild type SdeA or indicated SdeA mutants, after 24 hours cells were collected and lysed, the total cell lysates were separated with SDS-PAGE and blotted with antibody against GRASP55. (**D**) Modification of GRASP65 by exogenous SdeA in cells. HEK293T cells were transfected with wild type SdeA or indicated SdeA mutants, after 24 hours cells were collected and lysed, GFP-tagged GRASP65 proteins were isolated from cell lysate and separated with SDS-PAGE followed by blotting with antibody against GFP.

### *Legionella* infection causes PR-ubiquitination of GRASP55 and GRASP65

To check whether these Golgi proteins are PR-ubiquitinated upon *Legionella* infection, we immunoprecipitated GFP-tagged GRASP55 and GRASP65 from HEK293T cells infected with *Legionella* strains and analyzed them for PR-ubiquitination. The results showed that both GRASP55 and GRASP65 were ubiquitinated in a time-depending manner following *Legionella* infection (Fig. 4A, B). Moreover, the ubiquitination level was increased in cells infected with the *Legionella ΔdupA/B* mutant strain, indicating that more PR-ubiquitinated protein accumulated in the absence of the deubiquitinases (Fig. 4A, B). *Legionella* infection-induced GRASP55 and GRASP65 PR-ubiquitination was lost when cells were infected with a strain that lacks SidE family effectors (*ΔsidEs*) (Fig. 4C, D), thus, confirming that these effectors are essential for PR-ubiquitination of host substrate proteins. Similar results were obtained for GCP60 (Figure 4-figure supplement 1A). To further confirm that this detected ubiquitination is exclusively PR-ubiquitination caused by SidE family effectors directly, we incubated GRASP55 or GRASP65, isolated from infected HEK293T cells, with purified DupA. As expected, DupA was able to remove the ubiquitination of GRASP55 and GRASP65 induced by *Legionella* infection (Figure 4-figure supplement 1B, C). These data suggest that SdeA PR-ubiquitinates Golgi tethering proteins GRASP55 and GRASP65 during *Legionella* infection, further supporting our hypothesis that this modification has a directed function.

**Figure 4.**
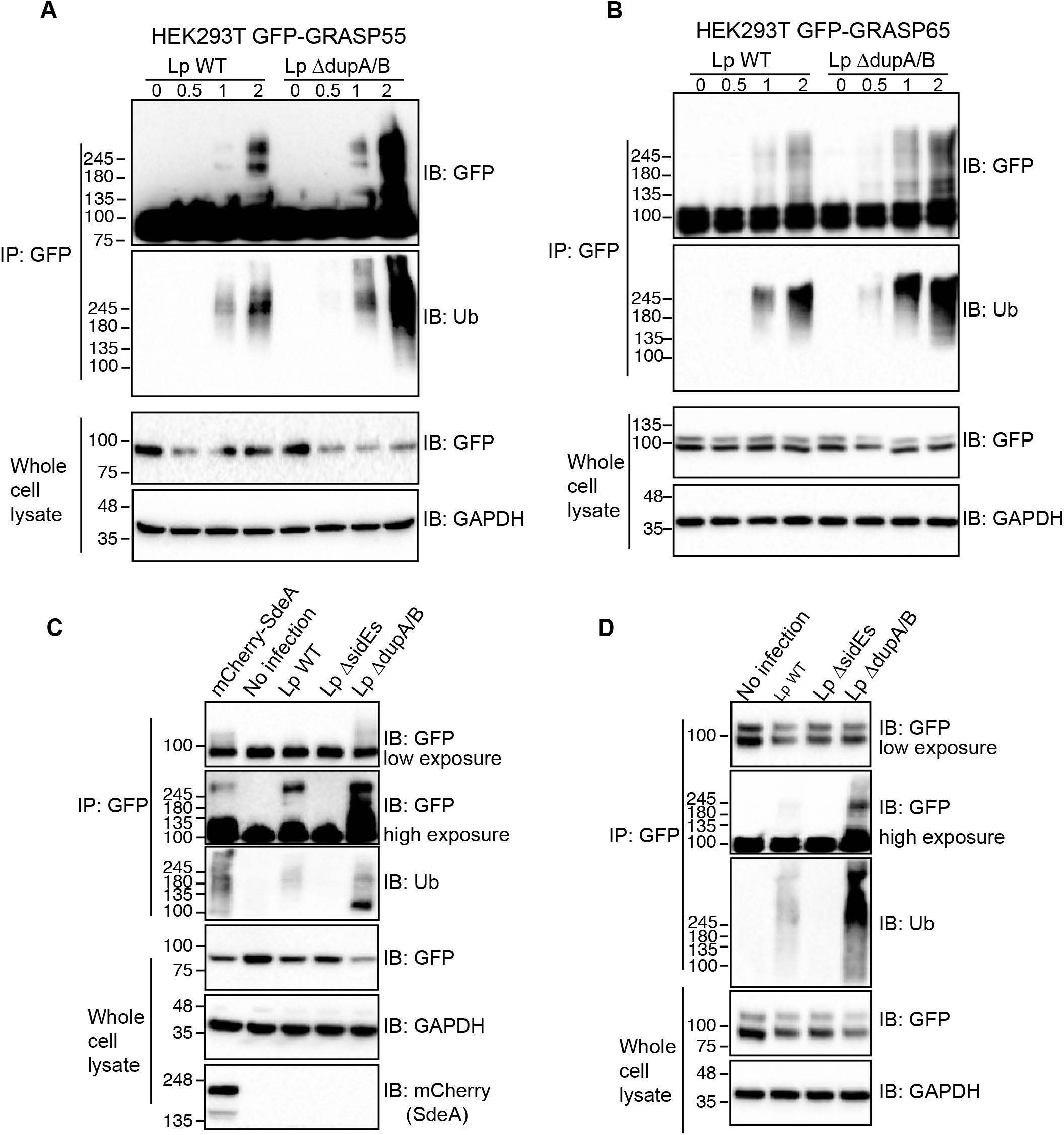
*Legionella* infection causes ubiquitination of GRASP proteins, which is dependent on SidE effectors. (**A**) Ubiquitination assay of GRASP55-GFP purified from HEK293T cells infected with *Legionella* strains. HEK293T cells were seeded in 6-well plate and co-transfected with plasmids encoding C-terminally GFP tagged GRASP55 and FcγRII then were infected for indicated times with *Legionella* bacteria opsonized by *Legionella* antibody. Cells were lysed with IP lysis buffer and purified GRASP55 proteins were separated by SDS-PAGE followed by blotting using anti-GFP and anti-ubiquitin antibodies. (**B**) Ubiquitination assay of GRASP65-GFP purified from HEK293T cells infected by *Legionella* wild type *or ΔdupA/B* mutant. (**C**) Ubiquitination assay of GRASP55-GFP purified from HEK293T cells infected with *Legionella* wild-type, *ΔsidEs or ΔdupA/B* strains. (**D**) Ubiquitination assay of GRASP65-GFP purified from HEK293T cells infected with *Legionella* wild-type, *ΔsidEs or ΔdupA/B* strains.

### SdeA ubiquitinates multiple serines of GRASP55 protein

Previous studies provided insights how SdeA targets and bridges Arg42 of Ub to serine residues of substrate proteins via a phosphoribosyl linker (Bhogaraju et al., 2016; Qiu et al., 2016). To gain insight into the mechanism of activity regulation of GRASP proteins by PR-ubiquitination, we used mass spectrometry to identify modified residues on GRASP55 following *in vitro* ubiquitination by SdeA (Fig. 5A). Four modified serine residues were identified in GRASP55 (S3, S408, S409, S449) (Fig. 5B, Figure 5-figure supplement 1). To further confirm these ubiquitination sites, we replaced seven serine residues (GRASP55 7S*), including the identified serines and their adjacent serines, by either threonine (S3, S4, S449, S451) or alanine (S408, S409, S441). We observed that ubiquitination of GRASP55 in cells co-expressing SdeA was markedly decreased when the serines were replaced (Fig. 5C). Similarly, we confirmed that GRASP55 bearing the seven mutated serine residues can not be ubiquitinated when cells were infected with wild-type or *ΔdupA/B Legionella* strains (Fig. 5D). These results confirm that these identified serines are the primary targets.

**Figure 5.**
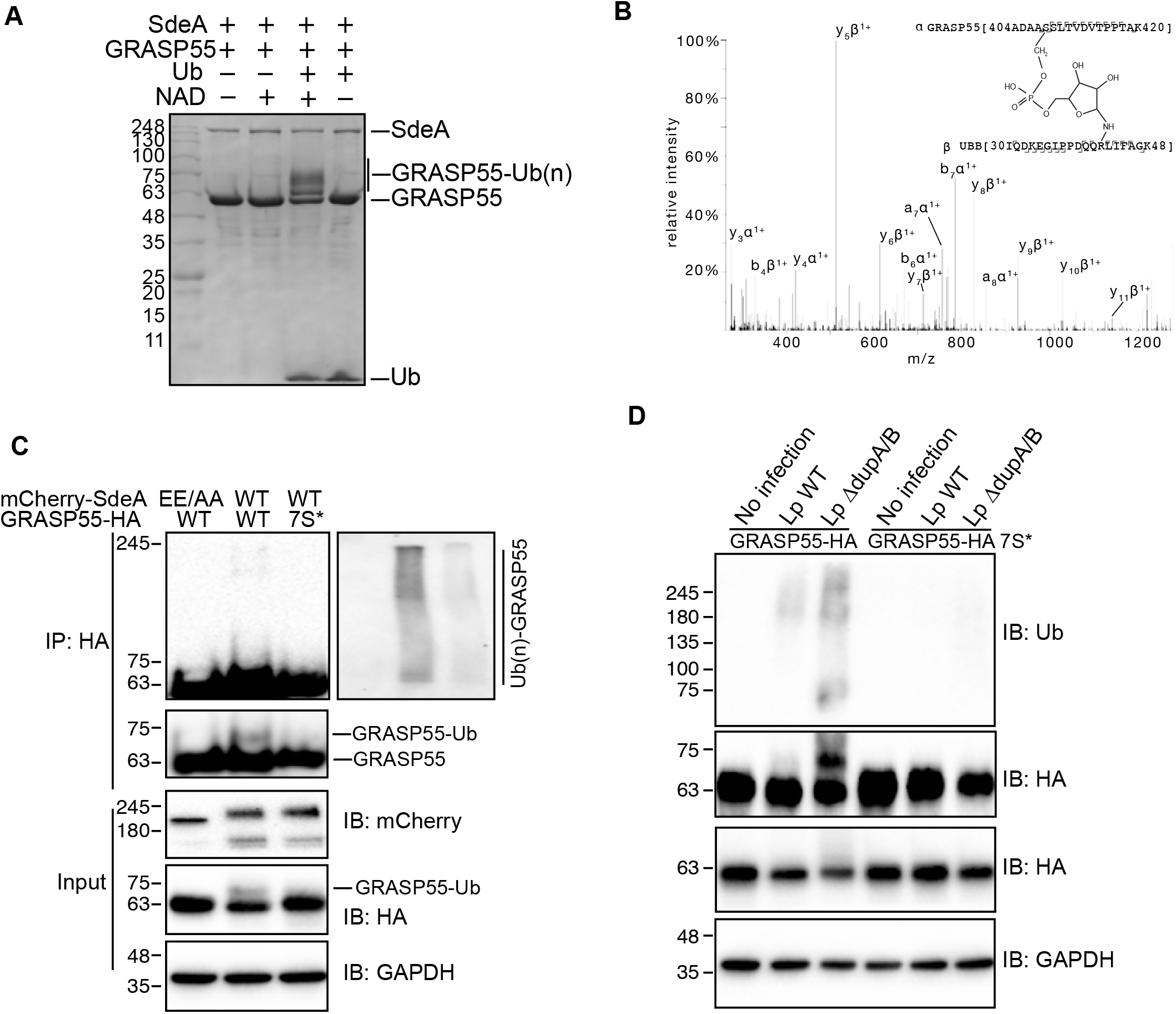
Identification of GRASP55 ubiquitination with mass spectrometry. (**A**) *In vitro* reaction of GRASP55 ubiquitination by SdeA for mass spectrometry analyses. 20 *μg* purified GRASP55 was incubated with SdeA and ubiquitin in the present of NAD. 10% reaction products were separated with SDS-PAGE and then stained with Coomassie blue or blotted with with antibodies against ubiquitin, GRASP55 to check the ubiquitination, the rest samples were subjected to mass spectrometry analyses. (**B**) Spectrum of GRASP S408-ubiquitin cross-linked peptide. (**C**) Validation of of ubiquitination sites in GRASP55. C-terminally HA-tagged wild-type and GRASP55 mutant were co-expressed with SdeA in HEK293T cells. After 24 h the cells were lysed for HA immunoprecipitation. Purified GRASP55-HA proteins were separated with SDS-PAGE followed by blotting using anti-HA and anti-ubiquitin antibodies. (**D**) Ubiquitination assay of wild type GRASP55 and mutant in cells infected with *Legionella*.

### PR-ubiquitination disrupts GRASP interactions

Studies have shown that GRASP proteins function in the connection of Golgi stacks and thereby Golgi structure maintenance through self-interaction and interactions with Golgi matrix proteins (Grond et al., 2020; Jarvela and Linstedt, 2012; Rabouille and Linstedt, 2016). Their activity can be regulated by post-translational modifications, for example, phosphorylation of serines within GRASP proteins was shown to result in Golgi fragmentation (Feinstein and Linstedt, 2008). We hypothesized that PR-ubiquitination of serines in GRASP proteins may affect self-interactions that are necessary for the connection of the Golgi stacks. To test this, we firstly PR-ubiquitinated purified GRASP55-GFP *in vitro* and then subsequently incubated the modified GRASP55 with purified His-tagged GRASP55. Co-IP analyses showed that PR-ubiquitinated GRASP55 exhibited reduced self-interaction compared to unmodified GRASP55 (Fig. 6A). This effect could also be seen in cells when the HA-tagged wild type GRASP55 or GRASP55 7S* serine mutant were co-expressed with GFP-tagged GRASP55 7S* in the presence of SdeA. The capacity of PR-ubiquitinated wild-type HA-GRASP55 to self-interact with GFP-GRASP55 7S*, was decreased in comparison to SdeA resistant HA-GRASP55 7S* (Fig. 6B). To analyze the functional impact of this observation on cells, we expressed wild-type GRASP55 or the GRASP55 7S* serine mutant in *GRASP55/GRASP65* KO HeLa cells, and then monitored the structural stability of the Golgi in cells co-expressing SdeA. As previously shown, double knockout of *GRASP55* and *GRASP65* induced dispersal of the Golgi (Bekier et al., 2017) (Figure 6-figure supplement 1). This phenotype could be rescued by ectopic expression of either wild-type GRASP55 or GRASP55 7S* (Figure 6-figure supplement 1), suggesting that serine mutations do not interfere with the function of GRASP55. Golgi disruption re-occurred when SdeA was concomitantly expressed with GRASP55 (Fig. 6C, D). However, the Golgi apparatus appeared less scattered when GRASP55 7S* was expressed, indicating that the higher resistance of GRASP55 serine mutant to SdeA ubiquitination activity results in increased structural stability of the Golgi in cells expressing SdeA (Fig. 6C, D). This data indicates that SdeA-caused Golgi disruption is supposedly the result of the modification of GRASP proteins, disturbing the connection between Golgi stacks.

**Figure 6.**
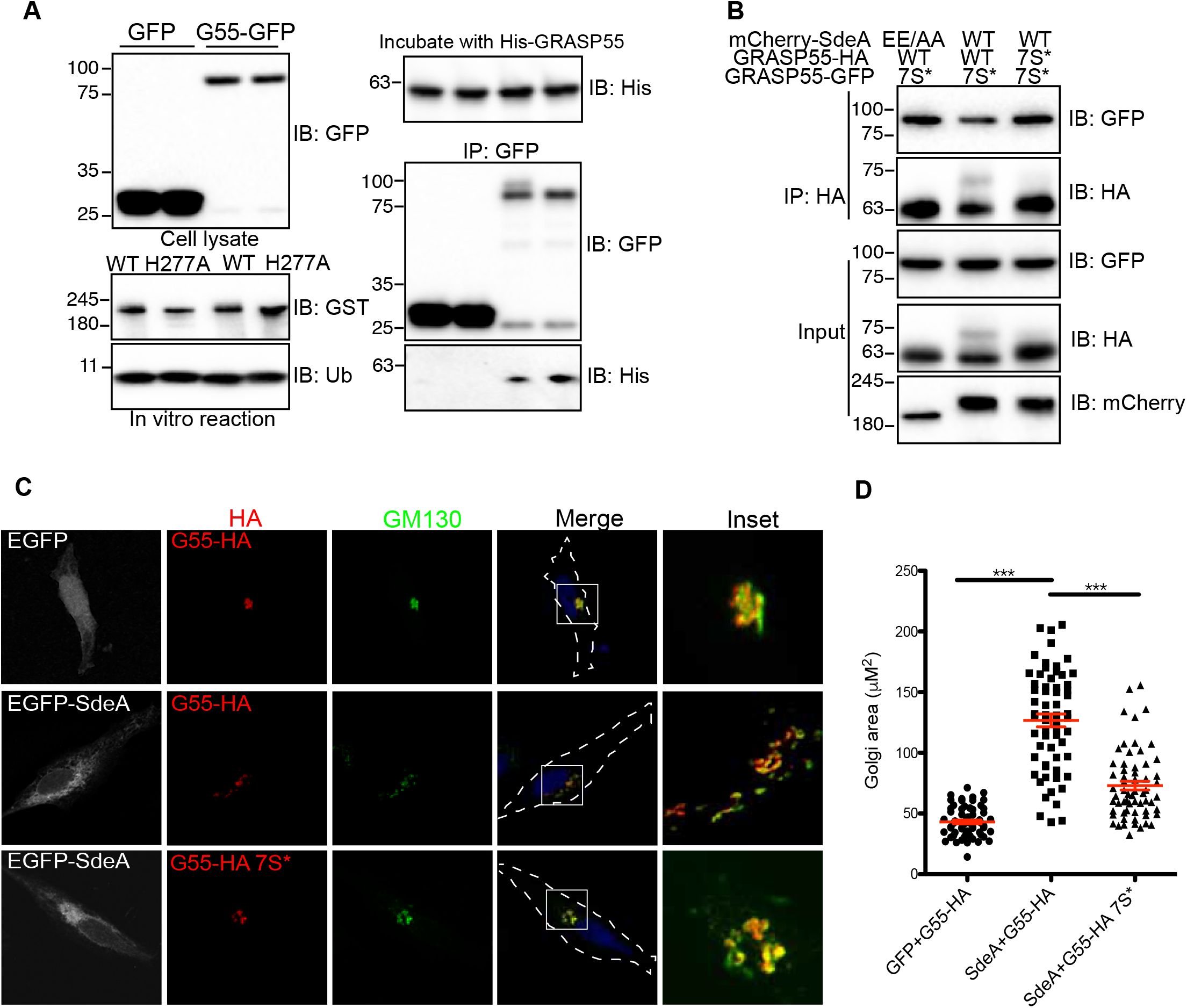
Serine ubiquitination impairs GRASP55 function. (**A**) Assay of the effect of PR-ubiquitination on GRASP55 dimerization *in vitro*. GRASP55-GFP purified from HEK293T cells were modified *in vitro* using SdeA and ubiquitin in the presence of NAD^+^, ubiquitinated GRASP55-GFP was then incubated with purified His-tagged GRASP55. Interaction between differently tagged GRASP55 proteins were analyzed with co-immunoprecipitation followed with western blotting. (**B**) Assay of the effect of PR-ubiquitination on GRASP55 dimerization *in vivo*. HA-tagged GRASP55 and GFP-tagged GRASP55 serine mutant were co-expressed with SdeA in HEK293T cells. Protein interaction between differently tagged GRASP55 were analyzed with co-IP and western blotting. (**C**) Confocal images showing that GRASP55 mutant is resistant to Golgi fragmentation caused by SdeA expression. Golgi areas of more than 60 cells from 3 replicates of each condition were measured with ImageJ software. (**D**) Data are shown as means ± SEM of more than 70 cells taken from three independent experiments. Data were analyzed with unpaired t test, ***P<0.001.

### *Legionella* containing vacuole does not recruit Golgi scatters

Intracellular pathogens tend to create a membrane surrounded niche for maturation, proliferation, and escape from defense mechanisms such as selective autophagy within the host cell. Along this line, *Chlamydia* infection induces Golgi fragmentation in order to generate Golgi ministacks for bacterial inclusions (Heuer et al., 2009). As for *Legionella, Legionella* containing vacuoles (LCVs) recruit ER membranes, thus converting the phagosome into a specific compartment that has features of ER (Kotewicz et al., 2017; Shin et al., 2020; Xu and Luo, 2013). We hypothesized that *Legionella* infection induces Golgi dispersal in order to facilitate the fusion of vesicles from the Golgi with LCV to enhance the formation of LCV and, ultimately, intracellular replication. To test this hypothesis, we infected HEK293T cells overexpressing GRASP55 or trans-Golgi marker GalT. The immunostaining showed that exogenous GRASP55 was recruited to LCV, however, our study recognizes the fact that exogenously overexpressed GRASP55 and GalT were shown to be partially localized in ER, which can be remodeled and recruited to LCV during infection. The recruited GRASP55 could very well be derived from the ER, and not the fragmented Golgi (Fig. 7A, B). To address whether *Legionella* indeed recruits fragmented Golgi cargo, we infected A549 cells with *Legionella*, stained cells with antibodies against endogenous cis-Golgi protein GM130 or trans-Golgi protein TGN46 and used microscopy to determine whether these Golgi markers are recruited to LCV upon infection. Immunostaining results suggested that neither cis-Golgi marker nor trans-Golgi accumulated on LCV (Fig. 7C, D). These data suggest that against our initial hypothesis *Legionella* does not disperse the Golgi simply to recruit Golgi-derived vesicles for the creation of LCVs, but that there must be another functional reasoning behind.

**Figure 7.**
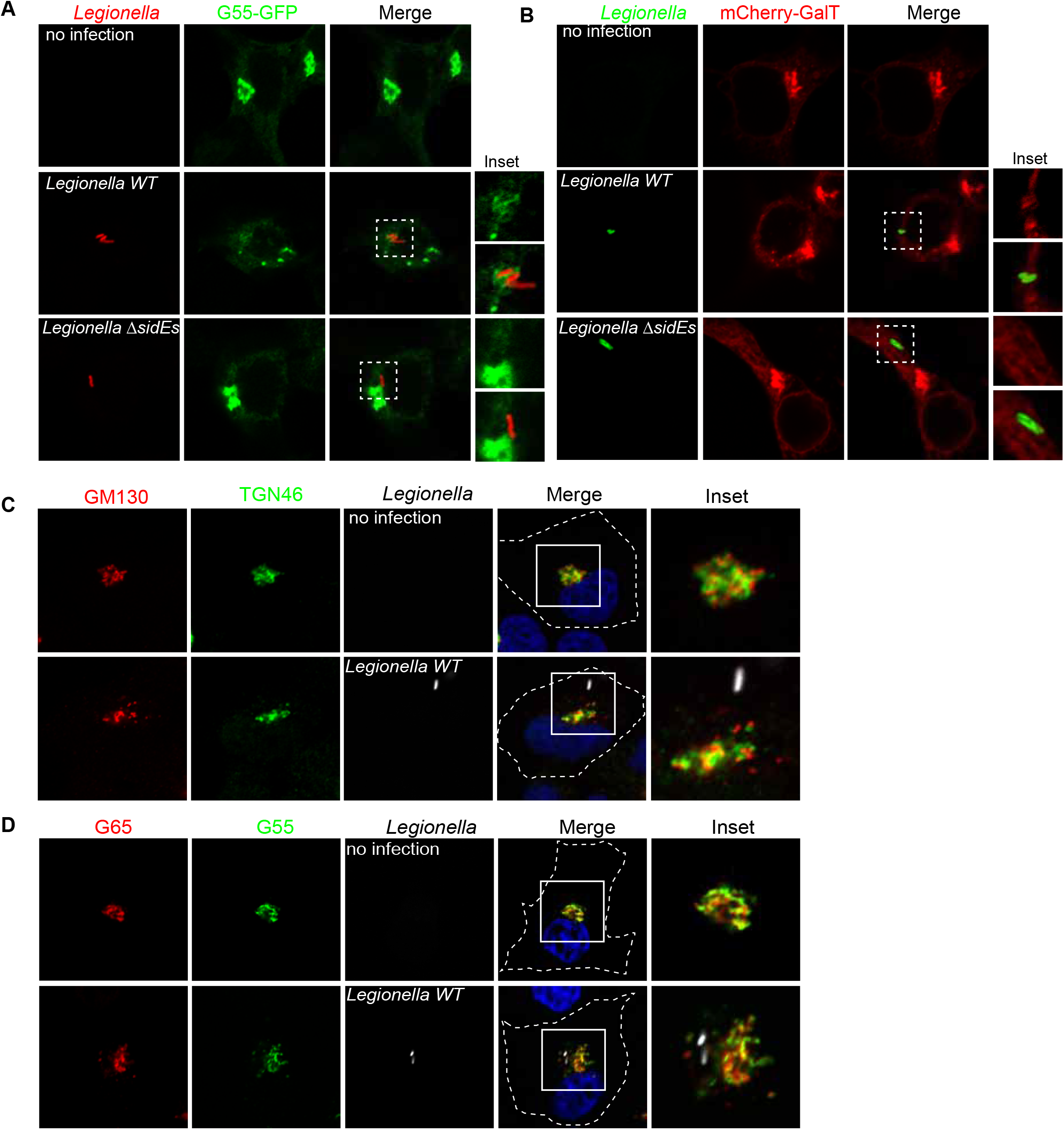
*Legionella* does not recruit fragmented Golgi. (**A**) Confocal images showing *Legionella* recruits overexpressed Golgi protein GRASP55. HEK293T cells transfected with plasmids encoding FCγRII and GFP-tagged GRASP55 were infected with indicated *Legionella* strains. Cells were washed 3 times with PBS after 2 hours infection to remove un-endocytosed bacteria, then fixed with 4% PFA and stained with antibody against *Legionella*. (**B**) Confocal images showing *Legionella* recruits overexpressed Golgi marker GalT. (**C**) Confocal images showing *Legionella* does not recruit endogenous cis-Golgi protein GM130 or trans-Golgi protein TGN46. A549 cells were infected with Legionella expressing dsRed and stained with antibodies against GM130 and TGN46.(**D**) Confocal images showing *Legionella* does not recruit endogenous cis-Golgi protein GRASP65 or trans-Golgi protein GRASP55. A549 cells were infected with Legionella expressing dsRed and stained with antibodies against GRASP65 and GRASP55.

### Serine ubiquitination regulates secretory pathway in host cells

In eukaryotic cells, the Golgi stack receives newly synthesized proteins from the ER, proteins then undergo modifications before being sorted via the trans-Golgi network. To investigate the effect of SdeA mediated Golgi disruption on Golgi function, we performed fluorescence recovery after photobleaching (FRAP) assay, the data indicates that the recovery of fluorescence after photobleaching of marked ROI is slower in SdeA expressing cells (Figure 8-figure supplement 1). Based on this observation, we then asked that whether SdeA inhibits protein trafficking via the Golgi. Vesicular stomatitis virus glycoprotein (VSVG) is a transmembrane protein that has been widely used as a tool to monitor protein trafficking through the secretory pathway (De Jong et al., 2006; Scidmore et al., 1996). This reporter contains a thermoreversible mutation which causes its misfolding and retention in the ER at 40 °C, while at the lower temperature of 32 °C, the protein folds correctly and is exported out of the ER to the plasma membrane via the secretory pathway (Bergmann, 1989; Presley et al., 1997). To access the functionality of the Golgi apparatus upon *Legionella* infection, we used a VSVG-GFP tracker protein to follow its transit through the Golgi, stained with the Golgi marker GM130. Immunofluorescence analyses indicated that in control A549 cells or cells infected with *Legionella* SidEs deletion strain, VSVG reached its peak of accumulation in the Golgi after 20 min of incubation at 32 °C, and the colocalization index then gradually decreased as the protein is trafficked from the Golgi to secretory vesicles. This process was slower in cells infected with wild-type *Legionella* or *ΔdupA/B* mutant strain, where maximal colocalization of VSVG with the GM130 occurred at a later time point and was more prolonged, indicating lower efficiency of protein trafficking through the secretory pathway (Fig. 8A, B). This was further confirmed by monitoring the sensitivity of VSVG glycosylation to Endoglycosidase H (EndoH). EndoH is an enzyme that removes mannose rich ER resident protein but not complex forms of N-like oligosaccharides from glycoproteins that are present in the Golgi or post Golgi compartments. The transformation of a glycoprotein from EndoH sensitive to EndoH resistant form has been widely used to monitor protein trafficking through the Golgi (Burke et al., 1984; Ernst et al., 2018). To specifically analyze the effect of PR-ubiquitination on VSVG trafficking through the Golgi with EndoH cleavage, we infected HEK293T cells at 40 °C and collected cells lysates at different time points after incubation at 32 °C, before treating them with EndoH. Western blots showed that VSVG trafficking was inhibited in cells infected with wild type *Legionella* or *Legionella* DupA/B deletion strain, compared with control cells or cells infected with *Legionella* ΔsidE strain. In control cells or cells infected with *Legionella* SidE deletion strain, the EndoH-resistant form of VSVG started to appear after 15 min incubation at 32 °C, and gradually accumulated over time, until almost all VSVG became EndoH-resistant form after 120 min. In cells infected with wild-type *Legionella*, the EndoH resistant-form of VSVG increased rather slowly, only around ~50% protein were converted to EndoH-resistant form at the same time point (Fig. 8C, D). These data further demonstrate that PR-ubiquitination caused by SidE effectors decelerates VSVG trafficking through the Golgi. This is also confirmed with a VSVG assay in cells expressing SdeA (Figure 8-figure supplement 2A, B). However, SdeA expression did not change the final glycosylation of LAMP1 in cells, as no significant band shift was detected on blot (Figure 8-figure supplement 2C). This suggests that activity of SdeA slows down trafficking through the Golgi but without completely inhibiting the function of the Golgi, this is in line with the fact that the SdeA does not impair the Golgi stacking.

**Figure 8.**
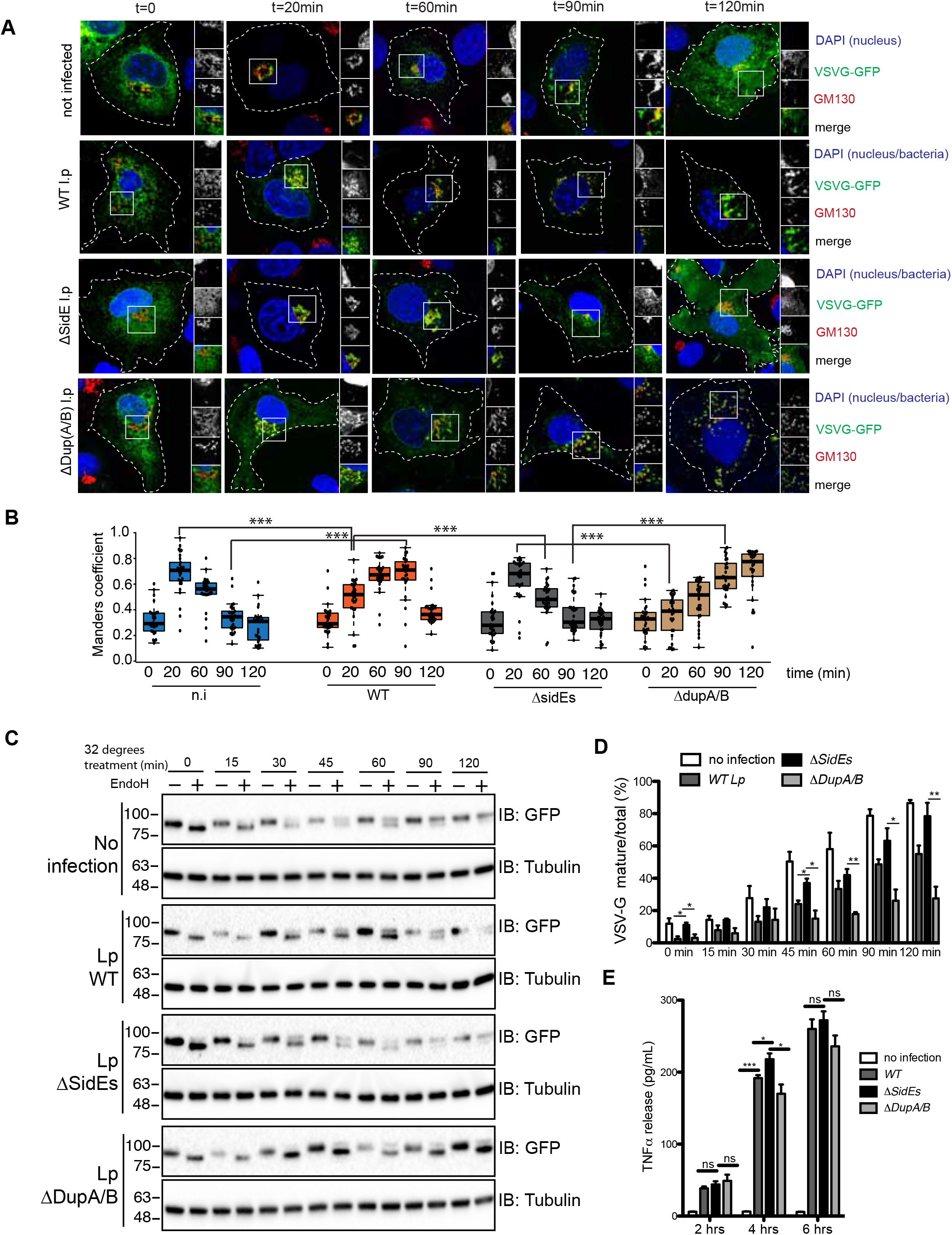
SdeA induced serine ubiquitination inhibits VSVG trafficking through Golgi membranes. (**A**) Confocal images showing the effect of SidE family effectors on VSVG trafficking during *Legionella* infection. (**B**) Quantitative analysis of the effect of SidE family effectors on VSVG trafficking during *Legionella* infection. Co-localization between VSVG and GM130 was shown as Manders coefficient. Data represents 30 cells taken from 3 independent experiments. White boxes indicate insets which are split into red, green, blue channels and displayed on the right side of the image. Center lines show the medians; box limits indicate the 25th and 75th percentiles as determined by R software; whiskers extend 1.5 times the interquartile range from the 25th and 75th percentiles, outliers are represented by dots; data points are plotted as circles. (**C**) Western blotting analysis of the effect of SidE family effectors on VSVG trafficking during *Legionella* infection using EndoH. Upper bands indicate the EndoH resistant form and lower bands indicate the EndoH sensitive form of VSVG. (**D**) Quantification of (**C**) to indicate the effect of *Legionella* infection on the conversion of EndoH sensitive form to resistant form of VSVG upon 32 °C incubation. Data were analyzed with unpaired t test, ***P<0.001, **P<0.01, *P<0.05. (**E**) ELISA assay of TNFα secreted from THP-1 cells infected with *Legionella* strains. Data are shown as means ± SEM of cytokine values of three independent experiments. Data were analyzed with unpaired t test, ***P<0.001, **P<0.01, *P<0.05.

As part of immune response, macrophage cells secret cytokines upon bacterial infection. Since ER-to-Golgi route trafficking plays an important role in conventional trafficking of most of the cytokines, and maintenance of Golgi structure is critical for secretion of some cytokines, such as TNFα (Micaroni et al., 2013), we decided to examine the effect of PR-ubiquitination on cytokine secretion of macrophage cells upon *Legionella* infection. THP-1 cells were treated with Phorbol 12-myristate 13-acxetate (PMA) to induce differentiation to macrophage cells, then cells were infected with wild type or *Legionella* strains. Media were collected and filtered for ELISA. The ELISA results show that cells infected with *Legionella* lacking SidE family effectors released more TNFα than cells infected with wild-type or *ΔdupA/B Legionella* strains (Fig. 8E). Taken together, these data demonstrate that Golgi disruption caused by SidE effectors impairs protein secretory pathway.

## Discussion

To date, considerable effort has been focused on investigating the mechanism and substrates of novel PR-ubiquitination catalyzed by SidE family of *Legionella* effectors. However, the functional consequences of PR-ubiquitination in the regulation of cellular processes has been poorly understood. In this study, we investigated the modification of Golgi proteins catalyzed by SidE effectors and explored the consequences of PR-ubiquitination in regulating Golgi morphology and secretory pathway.

The Golgi protein GRASP55 was identified as one of the most enriched candidates among all PR-Ub modified substrate proteins using mass-spectrometry (Shin et al., 2020). By conducting *in vitro* reactions and MS-based analyses, we identified several serine residues in GRASP55 potentially modified by SdeA. Mutation of the identified as well as adjacent serines markedly suppressed the PR-ubiquitination of GRASP55 both in cells expressing SdeA or infected with *Legionella*. Notably, mutation of these serines did not completely abolish the ubiquitination signal from purified GRASP55, suggesting that alternative residues in GRASP55 could also be modified by SdeA. This finding is consistent to other known substrates like Rab33b, in which S154 has been identified as a ubiquitination site for SdeA, yet S154A mutation does not abrogate ubiquitination (Bhogaraju et al., 2016). SdeA appears to modify substrate serine sites independent of specific structural motifs and serines in the flexible regions are prone to be modified as shown for Rab proteins (Wang et al., 2018).

Immunoblotting analyses of GRASP55 purified from cells either expressing SdeA or infected with *Legionella* revealed that PR-ubiquitinated GRASP55 is detected as high molecular weight smear under long exposure. This is likely due to the multi-ubiquitination event taking place on several serines of GRASP55, besides the identified preferred serines by SdeA. This hypothesis is supported by the observation that incubation with deubiquitinase DupA eliminated the high molecular bands from ubiquitinated GRASP proteins (Figure 4-figure supplement 1).

GRASP proteins contain a conserved N-terminal GRASP domain that is used to localize the proteins to the Golgi as well as to tether other GRASP proteins through trans-dimerization, which can be regulated through phosphorylation of the C-terminal serine and proline rich (SPR) domain by mitotic kinases (Feinstein and Linstedt, 2008; Wang et al., 2005). Several serines in this C-terminal region of GRASP55 including S408, S409, S441, S449 identified to be PR-ubiquitinated in this study, have also been reported to be phosphorylated in previous studies (Bian et al., 2014; Kim et al., 2016). Phosphorylation mimics at these sites disrupt the homodimerization of GRASP, possibly through protein conformational changes (Kim et al., 2016; Truschel et al., 2012). Based on the findings from a study using *in vivo* GRASP55/66 depletion, Grond et al. proposed that, instead of stacking the Golgi cisternae core, GRASP proteins function in linking of the rims of Golgi cisternae and the consequent connection of Golgi stacks (Grond et al., 2020). We show that PR-ubiquitination of GRASP55 also affects its homodimerization. Self-interaction of GRASP55 was diminished when the protein was PR-ubiquitinated by SdeA, both in *vitro* and *in vivo*. This disruption of GRASP protein homodimerization by PR-ubiquitination may lead to disconnection of Golgi stacks, but not complete fragmentation of the Golgi, which is supported by the confocal and super-resolution microscopy imaging of co-staining of cis and trans Golgi markers. Of note, in our previous study, some proteins related to, such as AKAP12, EPB41, SLK, were identified as possible substrates of SdeA (Shin et al., 2020). This may also affect the assembly of organelles including the Golgi. More studies will be needed in future to answer the question whether *Legionella* regulates cytoskeleton organization through SdeA mediated PR-ubiquitination of these cytoskeletal proteins.

Many pathogens have been characterized to require host organelles for their own intracellular survival and proliferation. As for *Legionella*, numerous host proteins have been detected on the LCVs. Of note, PI(4)P decoration on LCV, which functions to recruit bacterial effectors during infection, was shown to be derived directly from the Golgi body of host cells (Weber et al., 2018). However, our results indicate that *Legionella* does not recruit Golgi membrane pools containing endogenous GRASP55 or GRASP65, as these substrate proteins were not detected on the LCVs. Our data suggest that *Legionella* effectors disperse the Golgi but are not involved in the recruitment of Golgi components. This is in agreement with earlier studies, in which LCVs were purified from infected host cells and analyzed using proteomics approach, but Golgi proteins were rarely identified (Schmölders et al., 2017; Urwyler et al., 2009). During our ongoing study and preparation of this manuscript, Wan and colleagues reported Golgi fragmentation upon *Legionella* infection and the PR-ubiquitination of GRASP55. In their study it was shown that GRASP55 was recruited to LCV upon *Legionella* infection (Wan et al., 2019). However, it should be noted that, unlike the endogenous GRASP55 protein that mainly localizes to the Golgi apparatus, overexpressed GRASP55 in their study was detected as largely localized to ER that could be recruited to LCV. The recruited GRASP55 could very well be derived from the ER, but not the dispersed Golgi. Recently, a study reported that PI(4)P-containing vesicles derived from Golgi are involved in mitochondria division (Nagashima et al., 2020). Given the fact that mitochondrial dynamics is tightly modulated during *Legionella* infection (Escoll et al., 2017), it is possible that *Legionella* SdeA affects mitochondria fission to facilitate bacterial replication. Further efforts will be needed to address the effect of PR-ubiquitination mediated Golgi disruption on mitochondria.

The Golgi apparatus plays a central role in the secretory pathway. Using VSVG as a marker, we were able to dissect the effect of SidE mediated PR-ubiquitination on protein trafficking. Our findings provide insights into the functional roles of Golgi substrate PR-ubiquitination and subsequent Golgi disruption which impacts Golgi-associated protein secretory pathway. We have shown that PR-ubiquitination decelerates VSVG trafficking through the Golgi using microscopy and EndoH digestion assay. Moreover, we have found that secretion of cytokine TNFα was increased for THP-1 cells infected with *Legionella* lacking SidE family effectors, compared with cells infected with wild type *Legionella*. The opposite effect was observed in infection with *Legionella* missing DupA/B. This finding is consistent with previous study showing that SdeA expression inhibits secretion of secreted embryonic alkaline phosphatase reporter (SEAP) (Qiu et al., 2016).

Notably, multiple *Legionella* effectors have been suggested to regulate secretory pathways by yet unclear mechanisms (Machner and Isberg, 2006; Nagai et al., 2002). Identification of effectors involved in the regulation of the host secretory pathways will help us better understand both the bacterial pathogen and host cellular processes involved in infection, and thus further studies are needed.

Taken together, our study demonstrates that SdeA targets the Golgi and ubiquitinates Golgi tethering proteins GRASP55 and GRASP65, resulting in Golgi disruption and inhibition of secretory pathway. By revealing the biological consequences of PR-ubiquitination on Golgi proteins, our study provides a Golgi manipulation strategy, which *Legionella* utilizes to benefit bacterial infection and replication in host cells. It will be interesting to study whether PR-ubiquitination confers additional versatile mechanisms to facilitate bacterial infection by verifying more substrates of SidE effectors in future.

## Materials and Methods

### Antibodies and reagents

All reagents were from Sigma, Roche or Roth. The following antibodies were used: antibodies against HA (C29F4), GFP (sc-9996), GRASP65 (sc-374423), from Santa Cruz; ubiquitin (P4BD) and ubiquitin (ab7254) from Cell signaling and Abcam respectively; mCherry (ab125096), Tubulin (ab6046), Calnexin (ab22595), *Legionella* (ab20943) from abcam; GM130 (D6B1), GAPDH (D16H11) from Cell signaling; GM130 (610823) from BD for IF only; GRASP55 (10598-1-AP) from proteintech, TGN46 (AHP500) from Biorad. Monoclonal Anti-HA–Agarose antibody (HA-7) was purchased from Sigma.

### Cloning and mutagenesis

For protein expression in mammalian cells, GFP or mCherry tagged DupA, wild-type EGFP-SdeA and catalytically defective mutants SdeA H277A and SdeA EEAA were generated as described previously (Sagar Bhogaraju et al., 2016). SdeA plasmids were digested with BamHI and XhoI, then inserted into mCherry-C1 vectors digested with BamHI and XhoI to generate N terminally mCherry tagged wild-type and mutated SdeA. Deletion of SdeA was designed according to the known structure and sequence prediction analyses. Truncated deletions SdeA^1-972^ and SdeA^909-C^ were amplified from full-length SdeA cDNA and digested with BamHI and XhoI. The digested DNA fragments were inserted into pEGFP-C1 vectors digested with BamHI and XhoI. GFP or HA tagged GRASP55 and GRASP65-GFP were generated by PCR from GRASP55 or GRASP65 cDNA and digested with XhoI and BamHI or HindIII and KpnI respectively, then inserted into pEGFP-N1 or pHA-N1 vector. For generation of the GRASP55 7S* mutant, identified serines and adjacent serines S3, S4, S449, S451 were mutated to threonine to minimally effect the physio-chemical properties of these amino acids, in addition, S408, S409, S441 were mutated to alanine by site-directed mutagenesis. For protein expression in *E. coli*, SdeA was amplified from SdeA cDNA and digested with BamHI and XhoI. The digested DNA fragments were inserted into pGEX-6p-1 vector digested with BamHI and XhoI. GRASP55 and GRASP65 cDNA were amplified from mammalian vector and digested with NdeI and BamHI and cloned into pET15b and pGEX-6p-1 vector respectively. Serine to threonine or alanine mutations were generated by site-directed mutagenesis.

### Cell lines culture and Transfection

HEK293T, A549, COS7 cells were purchased from ATCC. Cells were cultured in high glucose Dulbecco’s Modified Eagles Medium (DMEM) supplemented with 10% fetal bovine serum (FBS), 100 U/mL penicilin and 100 mg/mL streptomycin at 37 °C, 5% CO_2_ in a humidified incubator. Transfection was performed using polyethyleneimine (PEI) reagent or Genejuice (Merck).

### *Legionella* culture and infection

*Legionella* strains were obtained from Dr. Zhao-Qing Luo lab (Purdue University). Cells were streaked and cultured at 37°C on N-(2-acetamido)-2-aminoethanesulfonic acid (ACES)-buffered charcoal-yeast extract (BCYE) agar plates for 3 days, followed by inoculation and growth for 20 h in 3 mL CYE liquid media. Post-exponential *Legionella* with OD_600_ between 3.6-3.8 were used to infect A549 or HEK293T cells. HEK293T cells were transfected with FCγRII and GRASP55-GFP or GRASP65-GFP for 24 hrs. Indicated *Legionella* strains were opsonized with antibody against *Legionella* (1: 500) at 37 °C for 30 min before infection. The HEK293T cells were infected with different *Legionella* strains at an MOI of 2 (for confocal imaging), or 10 (for Western blot) for the indicated time.

### SdeA mediated PR-ubiquitination reaction

SdeA mediated PR-Serine ubiquitination *in vitro* reaction was done as previously described (Kalayil et al., 2018). Briefly, 5 μM GRASP proteins were incubated with 1 μM of SdeA and 25 of μM ubiquitin in the presence of 200 μM of NAD^+^ in 40 μL of reaction buffer (50 mM NaCl and 50 mM Tris, pH 7.5) for 1 hour at 37 °C. Deubiquitination assay were performed by incubating PR-ubiquitinated proteins with 1 μg of GST-DupA at 37 °C for 1 h in reaction buffer (150 mM NaCl, 50 mM Tris-HCl pH 7.5). The reaction products were analyzed by SDS-PAGE with Coomassie staining or western blotting using antibodies against GST (cell signaling technology), His (Roche), GRASP55 (Proteintech), GRASP65 (Sino biotech.), Ub (Abcam, or Cell signaling technology). To assess the PR-ubiquitination of GRASP55 and GRASP65 in cells, plasmids for expression of GRASPs-GFP, GFP-SdeA or mCherry-SdeA, were co-transfected into HEK293T cells, cells were then cultured at 37 °C for 24 h. Whole cell lysates were subjected to immunoprecipitation with GFP-trap beads and the products or the whole cell lysates were separated with SDS-PAGE and blotted with antibodies against GFP or GRASP proteins.

### Western blotting and Immunoprecipitation

Cell lysates or immunoprecipitated proteins were mixed with SDS sample buffer, heated at 95 °C for 5 min, centrifuged, and separated by Tris-Glycine SDS-PAGE, and transferred to PVDF membrane (Millipore) at cold room. Blots were blocked with 5% nonfat milk for 1 hour at room temperature and incubated with primary antibodies overnight at cold room or 2 hours at room temperature and washed with TBST (0.1% Tween 20 in TBS) three times. The blots were further incubated with secondary antibodies for 1 h at room temperature and washed 3 times with TBST. The blots were incubated with ECL reagents (advansta), and chemiluminescence was acquired with the Bio-Rad ChemiDoc system. For immunoprecipitation, HEK293T cells expressing GFP or HA-tagged proteins were lysed with mild immunoprecipitation buffer containing 150 mM NaCl, 50 mM Tris-HCl, pH 7.5, 0.5% NP40, 1 mM PMSF, protease inhibitor cocktail (Sigma Aldrich), mixed with 10 μL GFP-trap or HA antibody conjugated agarose, and incubated for 4 h in cold room with end to end rotation. Beads were washed 3 times in IP buffer containing 500 mM NaCl. Proteins were eluted by resuspending with 2X SDS sample buffer followed by boiling for 5 min at 95 °C. Samples were then submitted to western blotting analysis.

### Protein expression and purification

GRASP55 and GRASP65 cDNA were cloned into p15b and pGEX-6p-1 vector respectively. Full-length SdeA was cloned into pGEX-6P-1 vector. *E. coli* competent cells (NEB T7 express) were transformed with plasmid, colonies were inoculated and cultured in LB medium overnight at 37 °C, The next day 5 mL culture was transferred to 1 L flask for further culture at 37 °C until the OD_600_ reaches to 0.6-0.8. Protein expression was induced by adding 0.5 mM IPTG and cells were further cultured overnight at 18 °C. The cells were harvested and the cell pellet was resuspended in lysis buffer (300 mM NaCl, 50 mM Tris-HCl pH 7.5) followed with sonication and centrifuged at 13,000 rpm to clarify the supernatant. Clarified lysates were then incubated with TALON beads or glutathione-S-Sepharose pre-equilibrated with washing buffer. Once eluted, proteins were further concentrated with filters and then purified by anion exchange chromatography on HitrapQ (GE Healthcare) and collected fractions were further loaded onto size exclusion column (Superdex 75 16/60, GE Healthcare). Proteins were concentrated and used for *in vitro* reaction.

### Identification of PR-ubiquitination serine sites on GRASP55

His-GRASP55 were purified from *E. coli* and PR-ubiquitinated by SdeA *in vitro*. Samples were prepared as previously described (Bhogaraju et al., 2016; Kalayil et al., 2018). Briefly, added urea buffer containing 8 M urea, 0.1 M Tris, pH 7.5 to the reaction mixture to a final volume of 200 μL, the reactions were then transferred to 30 kDa filter (Amicon Ultra, 0.5 mL, Merck) and washed 3 times with 200 μL of urea buffer by centrifugation to remove free ubiquitin. Proteins were washed 2 times with 50 mM ABC, pH 7.5, then digested with trypsin in 50 mM ABC pH 7.5 at trypsin to protein ratio 1:50 for 6 h and subsequently desalted by C18 and analyzed by LC MS/MS.

### Data quantification

Data shown in Figure 2B, 2D, 6D, 8B, 8D, 8E were analyzed with GraphPad Prism 5.0. Three independent experiments were performed, p values were determined using unpaired two-tailed t test, ***, **, * and ns represent p<0.0001, p<0.01, p<0.05 and not significant respectively. For Figure 2B and 2D, more than 70 SdeA transfected or *Legionella* infected cells were examined from 3 replicates of each condition. Values of percentage of cells with fragmented Golgi were input into GraphPad Prism, and analyzed. For Figure 6D, Golgi areas of more than 60 cells from 3 replicates of each condition were measured with ImageJ software. For Figure 8B, data represents 30 cells taken from 3 independent experiments. For Figure 8D, gray values of VSVG bands shown as Figure 8C from 3 replicates were measured with ImageJ.

### VSVG trafficking assay

HEK293T or A549 cells were co-transfected with VSVG-GFP and FcγRII or transfected with VSVG-GFP respectively and cultured at 37 °C for 24 h to express the proteins before being transferred to 40 °C. After 16 h incubation at 40 °C, cycloheximide was added into medium to inhibit further protein synthesis, after 2 h treatment cells were infected with *Legionella* for another 2 h then washed 3 times with PBS and cultured with fresh medium at 32 °C to remove the bacteria outside host cells, and then moved to 32 °C for different time points to release VSVG from ER. A549 cells were fixed and VSVG trafficking was acquired with confocal microscopy after immunofluorescence staining. DAPI marks nucleus and cytosolic bacteria. For calculating Manders coefficient in FIJI, ROIs of 30 μm^2^ are chosen from the perinuclear region containing the Golgi marked by GM130. Manders coefficient is calculated using Coloc2 plugin in FIJI and denotes fraction of VSVG-GFP pixels that is positive for GM130. For EndoH cleavage assay, HEK293T cells were lysed with lysis buffer containing 1% SDS, 50mM Tris, pH 8.0. Benzonase was added to reduce the viscosity caused by released DNA. Cell lysates were mixed with denaturing buffer then boiled for 10 min at 95 °C. Denatured proteins were incubated with EndoH for 3 h at 37 °C to cleave the EndoH sensitive form of glycosylation, final products were separated with SDS-PAGE and the EndoH-caused band shift was analyzed by blotting GFP.

### ELISA

To investigate the effect of PR-ubiquitination on secretion pathway, we analyzed the secretion of pro-inflammatory cytokine in PMA-treated THP-1 cells upon infection with *L*. *pneumophila*. Cytokine secretion analysis was performed with ELISA kit ordered from R&D system (TNFα: DY210-05) according to the manufacturer’s instruction.

### Immunofluorescence

HEK293T, COS7 or A549 cells were seeded on a coverslip in 12-well plates and cultured in CO_2_ incubator. Next day cells were transfected with plasmids encoding SdeA. The immunostaining was performed following the protocol previously described (Bhogaraju et al., 2016). Briefly, cells were washed once with PBS, pH 7.4, and fixed with 4% paraformaldehyde (PFA) in PBS for 10 min at room temperature. Cells were washed again with PBS 2 times, then permeabilized with 0.1% saponin in PBS for 10 min, and blocked with blocking buffer containing 0.1% saponin and 2% BSA in PBS for 1 h at room temperature. Cells were stained with antibodies diluted in blocking buffer overnight at 4 °C and washed with PBS three times next day. Cells were further incubated with Alexa Flour dyes-conjugated secondary antibodies for 1 h at room temperature in the dark and washed with PBS and incubated with DAPI in PBS, followed with further 2 times washing with PBS. Confocal imaging was performed using the Zeiss LSM780 microscope system. Images were analyzed with Fiji software.

### DNA-PAINT super-resolution Imaging

COS7 cells were fixed with prewarmed (37 °C) 4% methanol-free formaldehyde (Sigma-Aldrich, Germany) in PBS for 10 min followed by three washing steps with PBS. Cells were quenched with 0.1% NaBH4 (Sigma Aldrich, Germany) for 7 min in PBS and washed thrice with PBS. Fixed cells were permeabilized and blocked in permeabilization/blocking-buffer, followed by incubation of primary antibodies against GM130 and Golgin 97 in permeabilization/blocking buffer for 90 min. Washed cells were then incubated with DNA-labeled secondary antibodies for 1 h and washed again. Finally, samples were post-fixed using 4% methanol-free formaldehyde for 10 min at room-temperature followed by three washing steps with PBS. For sequential DNA-PAINT imaging, 125 nm gold-beads (Nanopartz, USA) were used as fiducial markers. Exchange DNA-PAINT measurements were performed at the N-STORM super-resolution microscopy system (Nikon, Japan) equipped with an oil immersion objective (Apo, 100x, NA 1.49) and an EMCCD camera (DU-897U-CS0-#BV, Andor Technology, Ireland).

## Acknowledgments

We thank Zhao-Qing Luo for the kind gift of *Legionella* strains (wild-type and *ΔsidEs*, Hubert Hilbi for the kind gift of *Legionella* expressing dsRed, Yanzhuang Wang for the kind gift of GRASP55/65 Knockout HeLa cell line; Mohit Misra, Anne-Claire Jacomin and Alexandra Stolz for discussion and critical reading of the manuscript. This project was funded by European Research Council (ERC) under the Europena Union’s Horizon 2020 research and innovation programme (I.D., grant agreement No 742720), and LOEWE Main Research Focus DynaMem of the German Federal State of Hesse (to I.D.). Work in Ivan Matic’s laboratory was funded by the Deutsche Forschungsgemeinschaft (DFG, German Research Foundation) under Germany’s Excellence Strategy-CECAD, EXC 2030-390661388 (to I.M.), and the EMBO Young Investigator Programme (to I.M.)

## Author contributions

Y.L. and I.D. designed the study and experiments. R.M. performed the VSVG trafficking experiment. F.B., T.C. and I.M. performed mass spectrometry experiments and data-analysis. Y.L. performed biochemical, cell biological and bacterial infection experiments and data-analysis. M.G. and M.H. performed DNA-PAINT super-resolution microscopy imaging. Y.L. and I.D. wrote the manuscript and all authors commented on it.

---

**Figure 2-figure supplement 1.**
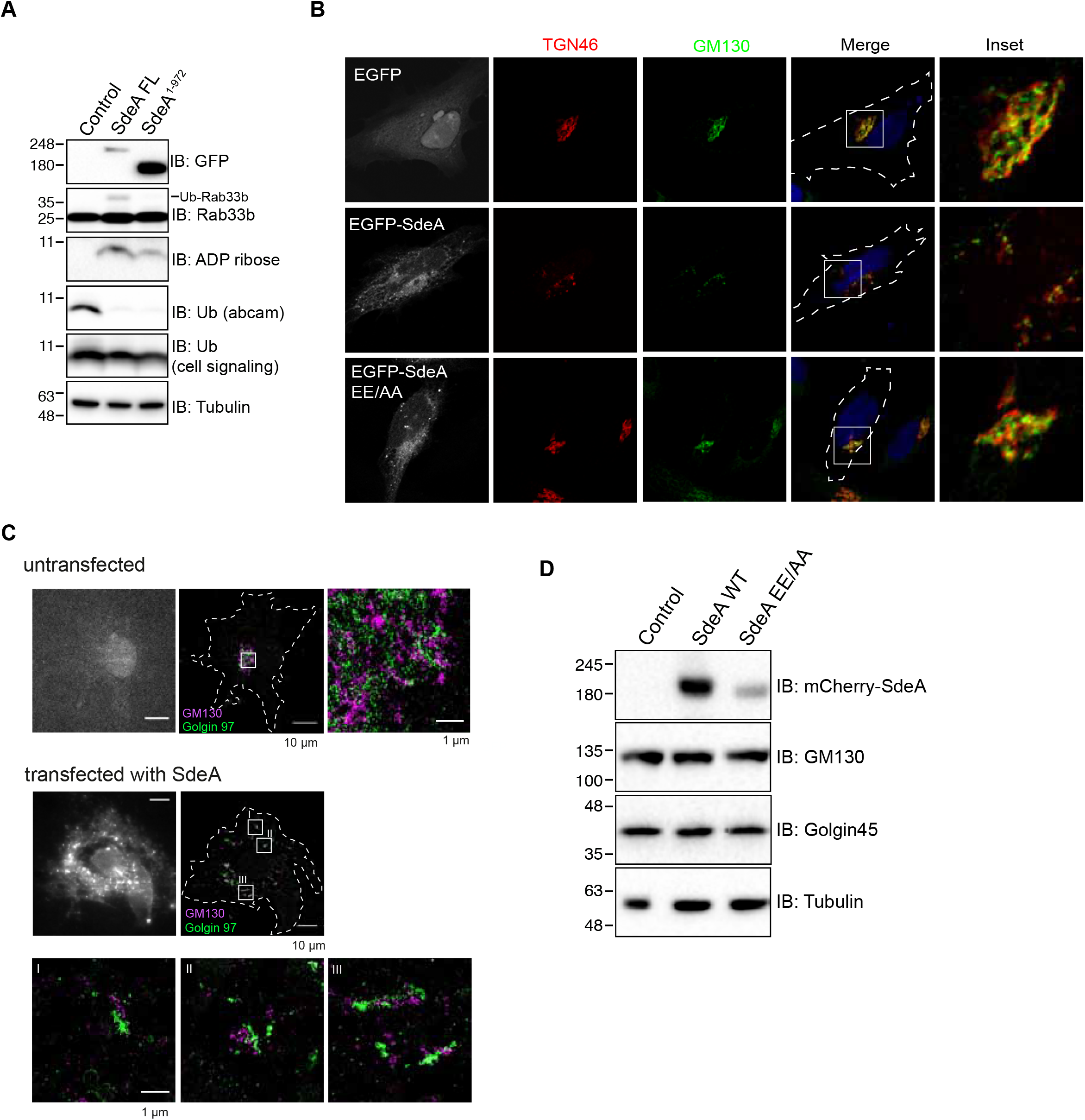
(**A**) Western blot analysis of modification of Golgi protein substrate by wild-type GFP-tagged SdeA or SdeA^1-972^ missing membrane targeting region. HEK293T cells were transfected with full-length or truncated SdeA, cells were lysed and blotted after 24 hours transfection. (**B**) Confocal images showing SdeA expression fragments the Golgi in HeLa cells. GFP-tagged SdeA wild type or catalytically defective mutants were expressed in HeLa cells. Cells were cultured for 24 hours after transfection then fixed with 4% PFA, after permeabilization, cells were stained with antibodies against cis-Golgi and trans-Golgi markers GM130 and TGN46. (**C**) DNA-PAINT super-resolution microscopy images of COS7 cells expressing SdeA. Fixed cells were incubated with primary antibodies against cis-Golgi and trans-Golgi markers GM130 and Golgin97, followed by incubation with secondary antibodies labeled with short oligonucleotide sequences. Data acquisition was performed with the N-STORM super-resolution microscopy system. (**D**) Western blot analysis of Golgi proteins GM130 and Golgin45 in cells expressing SdeA.

**Figure 3-figure supplement 1.**
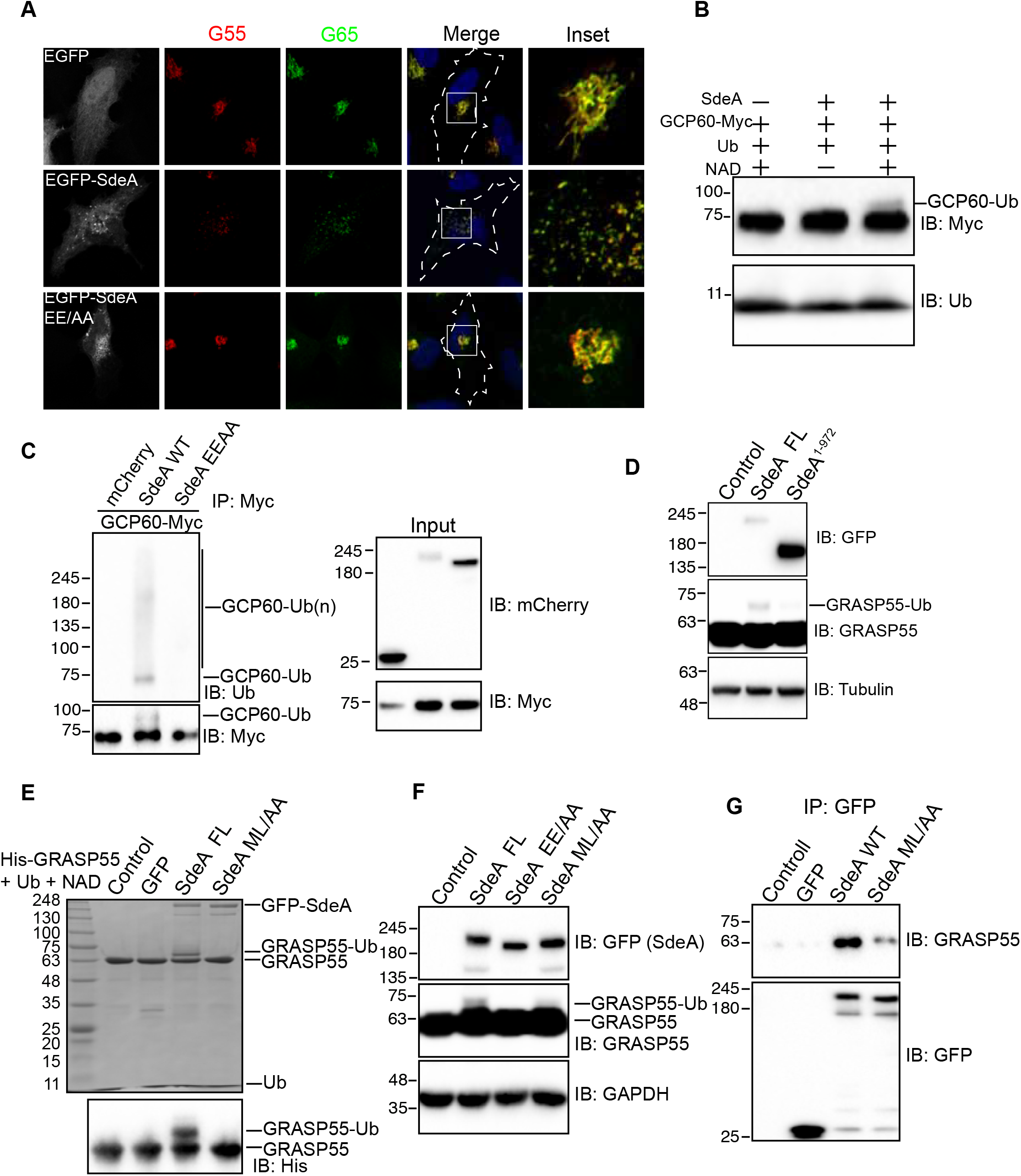
SdeA ubiquitinates Golgi tethering proteins. (**A**) Confocal images showing Golgi localization of endogenous GRASP55 and GRASP65. (**B**) GCP60 ubiquitination by SdeA *in vitro*. Purified Myc-tagged GCP60 was incubated with SdeA in the present of NAD^+^ and ubiquitin. Reaction products were blotted with antibodies against ubiquitin or Myc. (**C**) Modification of GCP60 by exogenous SdeA in cells. HEK293T cells were co-transfected with GCP60 and wild type SdeA or SdeA mutant, after 24 hours cells were collected and lysed, followed with Myc-IP. IP products were washed and separated with SDS-PAGE and blotted with antibody. (**D**) Modification of Golgi substrate GRASP55 by wild-type SdeA or SdeA^1-972^ missing membrane targeting region. (**E**) *In vitro* reaction of wild-type or ML/AA mutant with purified GRASP55. Reaction products were separated with SDS-PAGE and then stained with Coomassie blue. Ubiquitinated GRASP55 bands were indicated. (**F**) Analysis of the effect of SdeA ML/AA mutant on PR-ubiquitination of GRASP55 in cells. (**G**) Assay of protein interaction between SdeA and GRASP55. HEK293T cells were transfected with GFP-tagged wild-type SdeA or SdeA ML/AA mutant, after 24 hours cells were lysed and GFP fusion proteins were isolated with GFP-trap beads. Then the pulled-down proteins were separated with SDS-PAGE and blotted with antibody against GRASP55.

**Figure 4-figure supplement 1.**
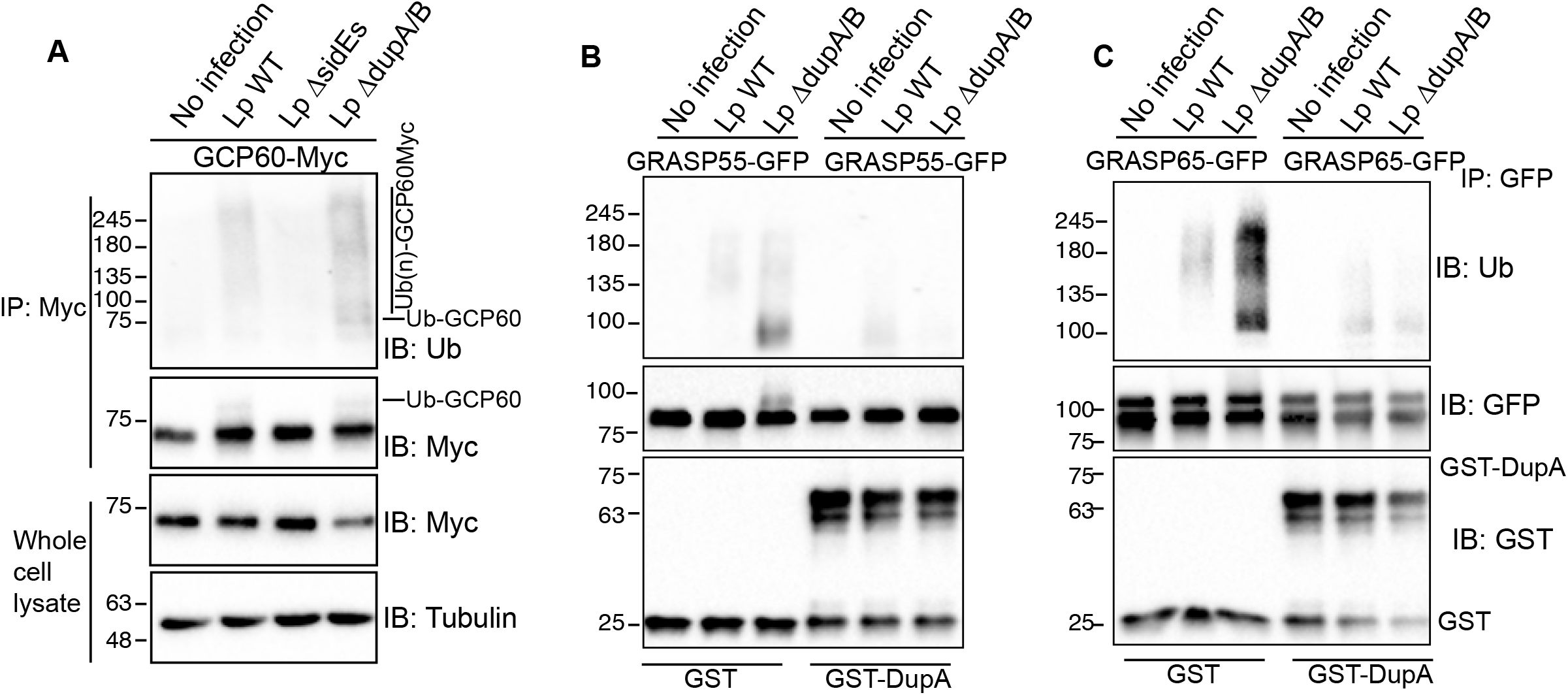
*Legionella* infection causes ubiquitination of Golgi proteins, which is dependent of SidE family proteins. (**A**) Ubiquitination assay of GCP60-Myc purified from HEK293T cells infected with *Legionella* strains. (**B**) Cleavage assay of PR-ubiquitination of GRASP55 with DupA. (**C**) Cleavage assay of PR-ubiquitination of GRASP65 with DupA.

**Figure 5-figure supplement 1.**
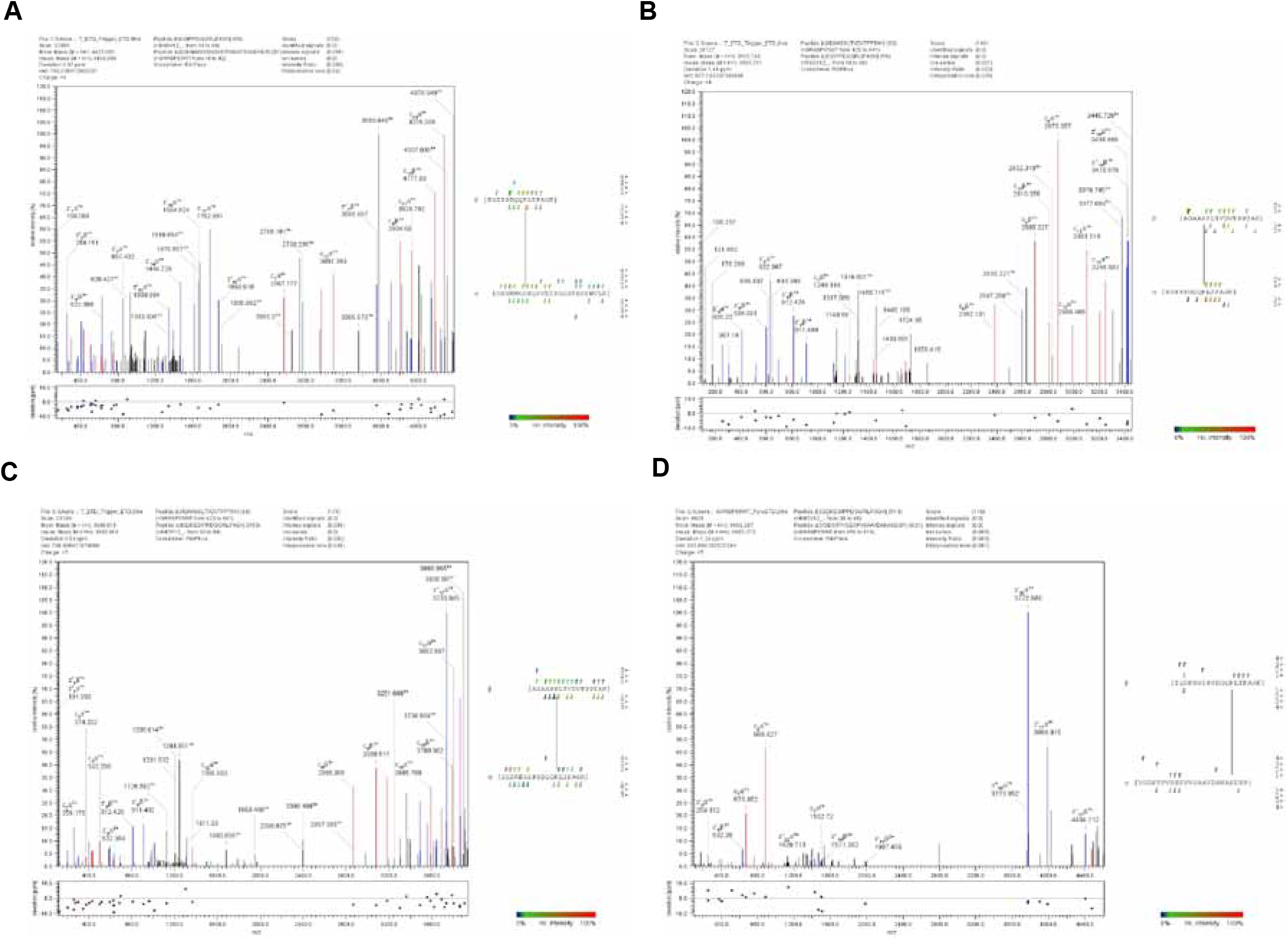
High resolution ETD spectrum of ubiquitin cross. (**A**) High resolution ETD spectrum of ubiquitin cross linked Serine 3 of GRASP55. (**B**) High resolution ETD spectrum of ubiquitin cross linked Serine 408 of GRASP55. (**C**) High resolution ETD spectrum of ubiquitin cross linked Serine 409 of GRASP55. (**D**) High resolution ETD spectrum of ubiquitin cross linked Serine 449 of GRASP55.

**Figure 6-figure supplement 1.**
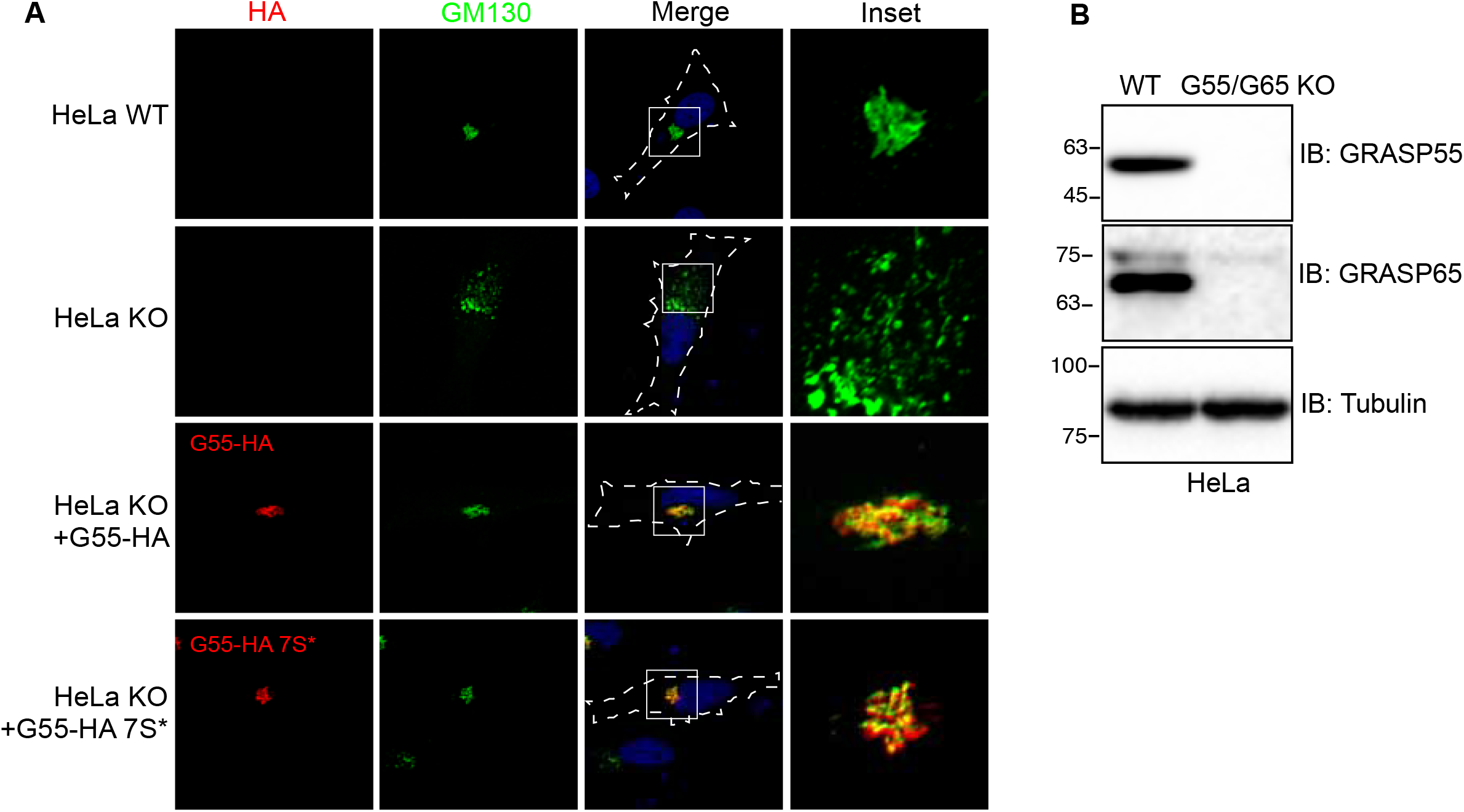
(**A**) Confocal images showing exogenously expressed wild-type GRASP55-HA and mutant rescue Golgi fragmentation caused by GRASP55/GRASP65 knockout. (**B**) Western blotting of cell lysates from wild-type and G55/G65 knockout HeLa cell lines. Knockout of G55 and G65 were validated with antibodies against G55, G65 respectively.

**Figure 8-figure supplement 1.**
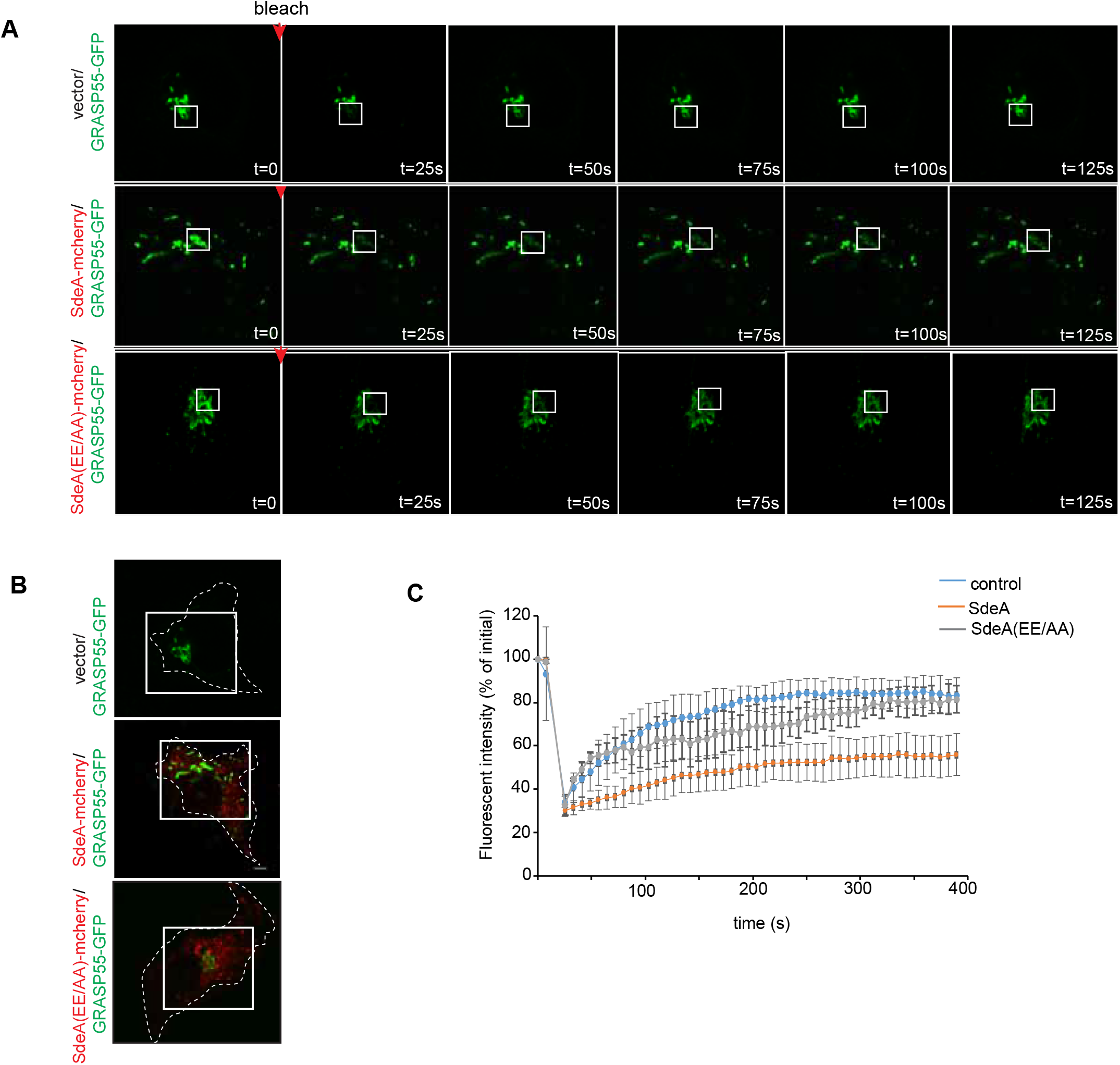
Golgi dynamics in cells expressing SdeA. (**A**) FRAP (fluorescence recovery after photobleaching) experiment showing Golgi dynamics in SdeA expressing cells. Golgi in live control or cells expressing SdeA (**B**) were photobleached at ROIs followed by measurement of fluorescence of ROIs over time. (**C**)Recovery curve of fluorescence after photobleaching of marked ROIs.

**Figure 8-figure supplement 2.**
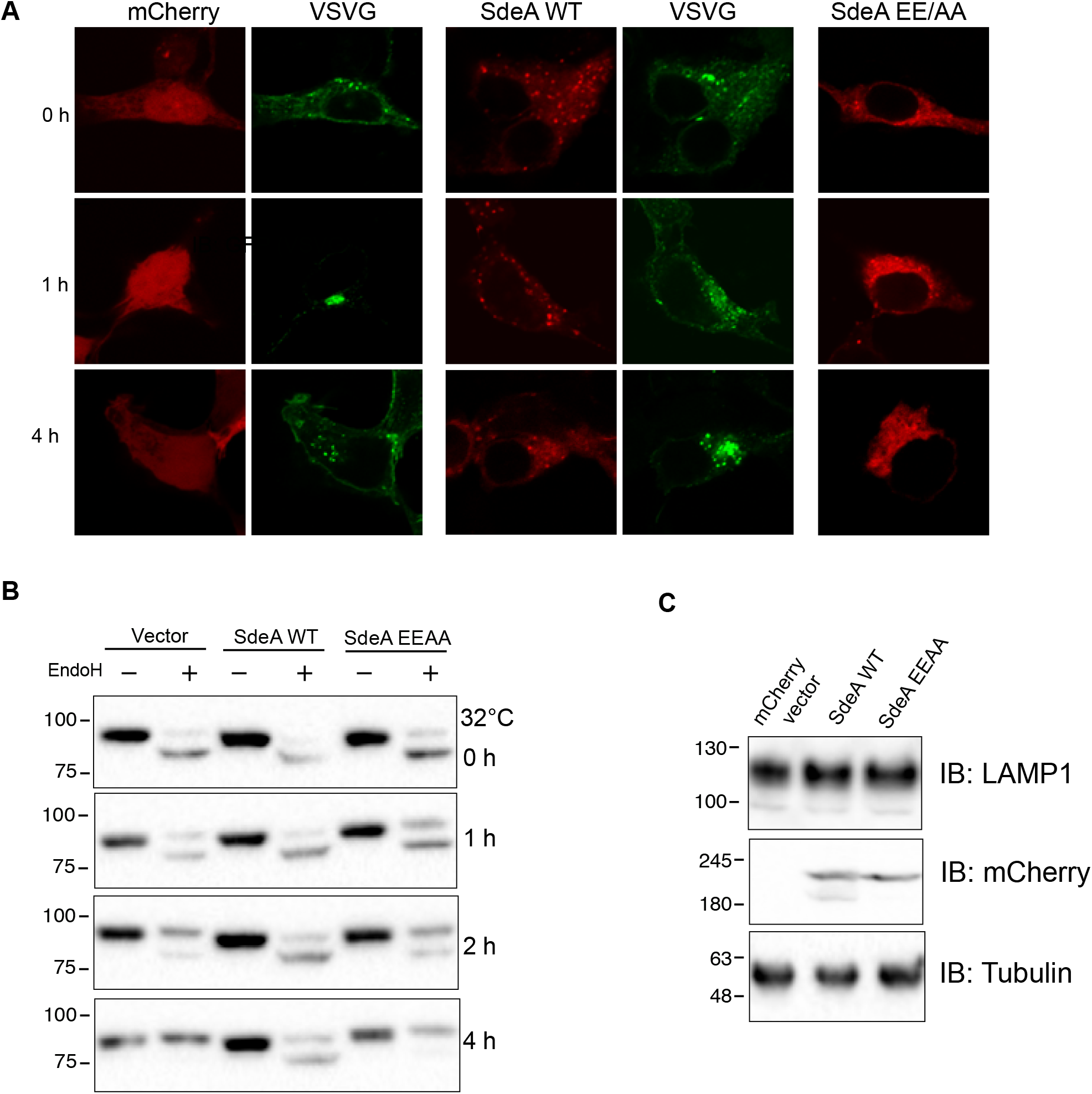
SdeA expression in cells impairs protein trafficking. (**A**) Analysis of VSVG trafficking in HEK293T cells expressing SdeA. Cells grown in 40 °C were moved to 32 °C for indicated time points. (**B**) Analysis of VSVG trafficking in HEK293T cells expressing SdeA using EndoH. Cells grown in 40 °C were moved to 32 °C for indicated time points. Cell lysates were probed with GFP antibody. (**C**) Analysis of the effect of SdeA expression on electrophoretic mobility of the Golgi protein LAMP1. Lysates from cells expressing SdeA WT or SdeA EEAA mutant were probed with LAMP1 antibody.

## References

Akturk A, Wasilko DJ, Wu X, Liu Y, Zhang Y, Qiu J, Luo ZQ, Reiter KH, Brzovic PS, Klevit RE, Mao Y. 2018. Mechanism of phosphoribosyl-ubiquitination mediated by a single legionella effector. Nature 557:729–745. doi: 10.1038/s41586-018-0147-6

Ashida H, Sasakawa C. 2016. Bacterial E3 ligase effectors exploit host ubiquitin systems. Curr Opin Microbiol 35:16–22. doi:10.1016/j.mib.2016.11.001

Bardill JP, Miller JL, Vogel JP. 2005. IcmS-dependent translocation of SdeA into macrophages by the Legionella pneumophila type IV secretion system. Mol Microbiol. doi:10.1111/j.1365-2958.2005.04539.x

Bekier ME, Wang L, Li J, Huang H, Tang D, Zhang X, Wang Y. 2017. Knockout of the Golgi stacking proteins GRASP55 and GRASP65 impairs Golgi structure and function. Mol Biol Cell 28:2833–2842. doi:10.1091/mbc.e17-02-0112

Ben-Neriah Y. 2002. Regulatory functions of ubiquitination in the immune system. Nat Immunol. doi:10.1038/ni0102-20

Bergmann JE. 1989. Using Temperature-Sensitive Mutants of VSV to Study Membrane Protein Biogenesis. Methods Cell Biol. doi:10.1016/S0091-679X(08)61168-1

Bhogaraju S, Kalayil S, Liu Y, Bonn F, Colby T, Matic I, Dikic I. 2016. Phosphoribosylation of Ubiquitin Promotes Serine Ubiquitination and Impairs Conventional Ubiquitination. Cell 167:1636–1649 e13. doi:10.1016/j.cell.2016.11.019

Bhogaraju Sagar, Kalayil S, Liu Y, Bonn F, Colby T, Matic I, Dikic I. 2016. Phosphoribosylation of Ubiquitin Promotes Serine Ubiquitination and Impairs Conventional Ubiquitination. Cell 167:1636–1649.e13. doi:10.1016/j.cell.2016.11.019

Bian Y, Song C, Cheng K, Dong M, Wang F, Huang J, Sun D, Wang L, Ye M, Zou H. 2014. An enzyme assisted RP-RPLC approach for in-depth analysis of human liver phosphoproteome. J Proteomics. doi:10.1016/j.jprot.2013.11.014

Bomberger JM, Ye S, MacEachran DP, Koeppen K, Barnaby RL, O’Toole GA, Stanton BA. 2011. A Pseudomonas aeruginosa toxin that hijacks the host ubiquitin proteolytic system. PLoS Pathog. doi:10.1371/journal.ppat.1001325

Burke B, Matlin K, Bause E, Legler G, Peyrieras N, Ploegh H. 1984. Inhibition of N-linked oligosaccharide trimming does not interfere with surface expression of certain integral membrane proteins. EMBO J. doi:10.1002/j.1460-2075.1984.tb01845.x

De Jong AS, Visch HJ, De Mattia F, Van Dommelen MM, Swarts HG, Luyten T, Callewaert G, Melchers WJ, Willems PH, Van Kuppeveld FJ. 2006. The coxsackievirus 2B protein increases efflux of ions from the endoplasmic reticulum and Golgi, thereby inhibiting protein trafficking through the Golgi. J Biol Chem. doi:10.1074/jbc.M511766200

Dikic I. 2017. Proteasomal and Autophagic Degradation Systems. Annu Rev Biochem. doi:10.1146/annurev-biochem-061516-044908

Donaldson KM, Yin H, Gekakis N, Supek F, Joazeiro CAP. 2003. Ubiquitin signals protein trafficking via interaction with a novel ubiquitin binding domain in the membrane fusion regulator, Vps9p. Curr Biol. doi:10.1016/S0960-9822(03)00043-5

Dong Y, Mu Y, Xie Y, Zhang Y, Han Y, Zhou Y, Wang W, Liu Z, Wu M, Wang H, Pan M, Xu N, Xu CQ, Yang M, Fan S, Deng H, Tan T, Liu X, Liu L, Li J, Wang J, Fang X, Feng Y. 2018. Structural basis of ubiquitin modification by the Legionella effector SdeA /631/80/458/582 /631/326/41/2536 /631/535 /631/45 /82/80 /82/83 /82/29 /82/16 /101/58 article. Nature 557:674–678. doi:10.1038/s41586-018-0146-7

Ernst AM, Syed SA, Zaki O, Bottanelli F, Zheng H, Hacke M, Xi Z, Rivera-Molina F, Graham M, Rebane AA, Björkholm P, Baddeley D, Toomre D, Pincet F, Rothman JE. 2018. S-Palmitoylation Sorts Membrane Cargo for Anterograde Transport in the Golgi. Dev Cell. doi:10.1016/j.devcel.2018.10.024

Escoll P, Song OR, Viana F, Steiner B, Lagache T, Olivo-Marin JC, Impens F, Brodin P, Hilbi H, Buchrieser C. 2017. Legionella pneumophila Modulates Mitochondrial Dynamics to Trigger Metabolic Repurposing of Infected Macrophages. Cell Host Microbe. doi:10.1016/j.chom.2017.07.020

Feinstein TN, Linstedt AD. 2008. GRASP55 regulates Golgi ribbon formation. Mol Biol Cell. doi:10.1091/mbc.E07-11-1200

Grond, R., Veenendaal, T., Duran, J. M., Raote, I., van Es, J. H., Corstjens, S., Delfgou, L., El Haddouti B.,, Malhotra, V., & Rabouille C. 2020. The function of GORASPs in Golgi apparatus organization in vivo. J Cell Biol. doi:10.1083/jcb.202004191

Hershko A, Ciechanover A, Varshavsky A. 2000. The ubiquitin system. Nat Med. doi: 10.1038/80384

Heuer D, Lipinski AR, Machuy N, Karlas A, Wehrens A, Siedler F, Brinkmann V, Meyer TF. 2009. Chlamydia causes fragmentation of the Golgi compartment to ensure reproduction. Nature 457:731–735. doi:10.1038/nature07578

Hicks SW, Galán JE. 2013. Exploitation of eukaryotic subcellular targeting mechanisms by bacterial effectors. Nat Rev Microbiol. doi:10.1038/nrmicro3009

Hubber A, Roy CR. 2010. Modulation of Host Cell Function by Legionella pneumophila Type IV Effectors. Annu Rev Cell Dev Biol. doi:10.1146/annurev-cellbio-100109-104034

Jarvela T, Linstedt AD. 2012. Golgi GRASPs: Moonlighting membrane tethers. Cell Health Cytoskelet. doi:10.2147/CHC.S21849

Jeong KC, Sexton JA, Vogel JP. 2015. Spatiotemporal Regulation of a Legionella pneumophila T4SS Substrate by the Metaeffector SidJ. PLoS Pathog. doi:10.1371/journal.ppat.1004695

Jubelin G, Frédéric T, Duda DM, Hsu Y, Samba-Louaka A, Nobe R, Penary M, Watrin C, Nougayréde JP, Schulman BA, Stebbins CE, Oswald E. 2010. Pathogenic bacteria target NEDD8-conjugated cullins to hijack host-cell signaling pathways. PLoS Pathog. doi:10.1371/journal.ppat.1001128

Kalayil S, Bhogaraju S, Bonn F, Shin D, Liu Y, Gan N, Basquin J, Grumati P, Luo ZQ, Dikic I. 2018. Insights into catalysis and function of phosphoribosyl-linked serine ubiquitination. Nature 557:734–738. doi:10.1038/s41586-018-0145-8

Kim J, Noh SH, Piao H, Kim DH, Kim K, Cha JS, Chung WY, Cho HS, Kim JY, Lee MG. 2016. Monomerization and ER Relocalization of GRASP Is a Requisite for Unconventional Secretion of CFTR. Traffic. doi:10.1111/tra.12403

Knodler LA, Ibarra JA, Pérez-Rueda E, Yip CK, Steele-Mortimer O. 2011. Coiled-coil domains enhance the membrane association of Salmonella type III effectors. Cell Microbiol. doi:10.1111/j.1462-5822.2011.01635.x

Kotewicz KM, Ramabhadran V, Sjoblom N, Vogel JP, Haenssler E, Zhang M, Behringer J, Scheck RA, Isberg RR. 2017. A Single Legionella Effector Catalyzes a Multistep Ubiquitination Pathway to Rearrange Tubular Endoplasmic Reticulum for Replication. Cell Host Microbe 21:169–181. doi:10.1016/j.chom.2016.12.007

Machner MP, Isberg RR. 2006. Targeting of Host Rab GTPase Function by the Intravacuolar Pathogen Legionella pneumophila. Dev Cell. doi:10.1016/j.devcel.2006.05.013

Maculins T, Fiskin E, Bhogaraju S, Dikic I. 2016. Bacteria-host relationship: ubiquitin ligases as weapons of invasion. Cell Res 26:499–510. doi:10.1038/cr.2016.30

Micaroni M, Stanley AC, Khromykh T, Venturato J, Wong CXF, Lim JP, Marsh BJ, Storrie B, Gleeson PA, Stow JL. 2013. Rab6a/a’ Are Important Golgi Regulators of Pro-Inflammatory TNF Secretion in Macrophages. PLoS One 8. doi:10.1371/journal.pone.0057034

Nagai H, Kagan JC, Zhu X, Kahn RA, Roy CR. 2002. A bacterial guanine nucleotide exchange factor activates ARF on Legionella phagosomes. Science (80-). doi:10.1126/science.1067025

Nagashima S, Tábara LC, Tilokani L, Paupe V, Anand H, Pogson JH, Zunino R, McBride HM, Prudent J. 2020. Golgi-derived PI(4)P-containing vesicles drive late steps of mitochondrial division. Science (80-). doi:10.1126/science.aax6089

Presley JF, Cole NB, Schroer TA, Hirschberg K, Zaal KJM, Lippincott-Schwartz J. 1997. ER-to-Golgi transport visualized in living cells. Nature. doi:10.1038/38001

Qiu J, Luo ZQ. 2017. Legionella and Coxiella effectors: strength in diversity and activity. Nat Rev Microbiol 15:591–605. doi:10.1038/nrmicro.2017.67

Qiu J, Sheedlo MJ, Yu K, Tan Y, Nakayasu ES, Das C, Liu X, Luo ZQ. 2016. Ubiquitination independent of E1 and E2 enzymes by bacterial effectors. Nature 533:120–124. doi:10.1038/nature17657

Qiu J, Yu K, Fei X, Liu Y, Nakayasu ES, Piehowski PD, Shaw JB, Puvar K, Das C, Liu X, Luo ZQ. 2017. A unique deubiquitinase that deconjugates phosphoribosyl-linked protein ubiquitination. Cell Res. doi:10.1038/cr.2017.66

Rabouille C, Linstedt AD. 2016. GRASP: A multitasking tether. Front Cell Dev Biol 4:1–8. doi:10.3389/fcell.2016.00001

Rape M. 2018. Ubiquitylation at the crossroads of development and disease. Nat Rev Mol Cell Biol 19:59–70. doi:10.1038/nrm.2017.83

Schmölders J, Manske C, Otto A, Hoffmann C, Steiner B, Welin A, Becher D, Hilbi H. 2017. Comparative proteomics of purified pathogen vacuoles correlates intracellular replication of Legionella pneumophila with the small GTPase ras-related protein 1 (Rap1). Mol Cell Proteomics. doi:10.1074/mcp.M116.063453

Scidmore MA, Fischer ER, Hackstadt T. 1996. Sphingolipids and glycoproteins are differentially trafficked to the Chlamydia trachomatis inclusion. J Cell Biol. doi:10.1083/jcb.134.2.363

Shin D, Mukherjee R, Liu Yaobin, Gonzalez A, Bonn F, Liu Yan, Rogov V V., Heinz M, Stolz A, Hummer G, Dötsch V, Luo ZQ, Bhogaraju S, Dikic I. 2020. Regulation of Phosphoribosyl-Linked Serine Ubiquitination by Deubiquitinases DupA and DupB. Mol Cell. doi:10.1016/j.molcel.2019.10.019

Shorter J, Watson R, Giannakou ME, Clarke M, Warren G, Barr FA. 1999. GRASP55, a second mammalian GRASP protein involved in the stacking of Golgi cisternae in a cell-free system. EMBO J. doi:10.1093/emboj/18.18.4949

Sohda M, Misumi Y, Yamamoto A, Yano A, Nakamura N, Ikehara Y. 2001. Identification and Characterization of a Novel Golgi Protein, GCP60, that Interacts with the Integral Membrane Protein Giantin. J Biol Chem. doi:10.1074/jbc.M108961200

Truschel ST, Zhang M, Bachert C, Macbeth MR, Linstedt AD. 2012. Allosteric regulation of GRASP protein-dependent golgi membrane tethering by mitotic phosphorylation. J Biol Chem. doi:10.1074/jbc.M111.326256

Urwyler S, Nyfeler Y, Ragaz C, Lee H, Mueller LN, Aebersold R, Hilbi H. 2009. Proteome analysis of Legionella vacuoles purified by magnetic immunoseparation reveals secretory and endosomal GTPases. Traffic. doi:10.1111/j.1600-0854.2008.00851.x

Wan M, Sulpizio AG, Akturk A, Beck WHJ, Lanz M, Faça VM, Smolka MB, Vogel JP, Mao Y. 2019. Deubiquitination of phosphoribosyl-ubiquitin conjugates by phosphodiesterase-domain–containing Legionella effectors. Proc Natl Acad Sci U S A. doi:10.1073/pnas.1916287116

Wang Y, Satoh A, Warren G. 2005. Mapping the functional domains of the Golgi stacking factor GRASP65. J Biol Chem. doi:10.1074/jbc.M412407200

Wang Y, Shi M, Feng H, Zhu Y, Liu S, Gao A, Gao P. 2018. Structural Insights into Non-canonical Ubiquitination Catalyzed by SidE. Cell 173:1231–1243.e16. doi:10.1016/j.cell.2018.04.023

Weber S, Steiner B, Welin A, Hilbi H. 2018. Legionella-Containing Vacuoles Capture PtdIns(4)P-Rich Vesicles Derived from the Golgi Apparatus. MBio 9. doi:10.1128/mBio.02420-18

Xu L, Luo ZQ. 2013. Cell biology of infection by Legionella pneumophila. Microbes Infect. doi:10.1016/j.micinf.2012.11.001

